# What drives chorismate mutase to top performance? Insights from a combined *in silico* and *in vitro* study

**DOI:** 10.1101/2022.11.08.515563

**Authors:** Helen V. Thorbjørnsrud, Luca Bressan, Tamjidmaa Khatanbaatar, Manuel Carrer, Kathrin Würth-Roderer, Gabriele Cordara, Peter Kast, Michele Cascella, Ute Krengel

## Abstract

Unlike typical chorismate mutases, the enzyme from *Mycobacterium tuberculosis* (MtCM) has only low activity on its own. Remarkably, its catalytic efficiency *k*_cat_/*K*_m_ can be boosted more than 100-fold by complex formation with a partner enzyme. Recently, an autonomously fully active MtCM variant was generated using directed evolution, and its structure solved by X-ray crystallography. However, key residues were involved in crystal contacts, challenging the functional interpretation of the structural changes. Here, we address these challenges by microsecond molecular dynamics simulations, followed up by additional kinetic and structural analyses of selected sets of specifically engineered enzyme variants. A comparison of wild-type MtCM with naturally and artificially activated MtCMs revealed the overall dynamic profiles of these enzymes as well as key interactions between the C-terminus and the active site loop. In the artificially evolved variant of this model enzyme, this loop is pre-organized and stabilized by Pro52 and Asp55, two highly conserved residues in typical, highly active chorismate mutases. Asp55 stretches across the active site and helps to appropriately position active site residues Arg18 and Arg46 for catalysis. The role of Asp55 can be taken over by another acidic residue, if introduced at position 88 close to the C-terminus of MtCM, as suggested by MD simulations and confirmed by kinetic investigations of engineered variants.

## INTRODUCTION

Pericyclic reactions are common in industrial processes, but very rare in biology (1–4). Chorismate mutase (CM) catalyzes the only known pericyclic process in primary metabolism, the Claisen rearrangement of chorismate (**1**) to prephenate (**2**), *via* a chair-like transition state (Scheme 1) (5). This catalytic step at the branch point of the shikimate pathway funnels the key metabolite chorismate towards the synthesis of tyrosine and phenylalanine, as opposed to tryptophan and several aromatic vitamins (6,7). The CM reaction is a concerted unimolecular transformation that is well studied both by experimental and computational means (8). It proceeds ostensibly *via* the same transition state both in solution and in enzyme catalysis (9,10). Due to these factors, CM has long been a model enzyme for computational chemists (11).

**Scheme 1.**
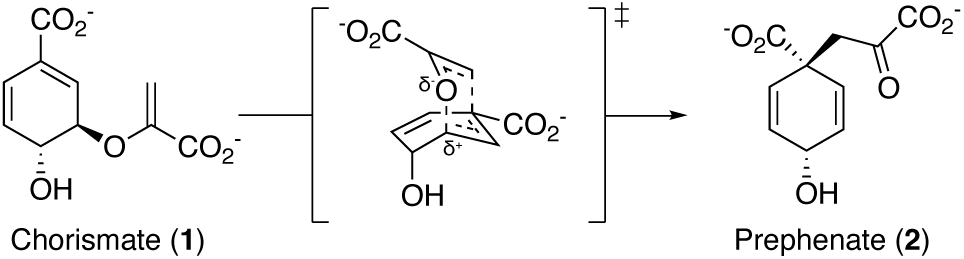
Chorismate mutase reaction. The Claisen rearrangement catalyzed by chorismate mutase converts chorismate (**1**) to prephenate (**2**) and proceeds *via* a highly polarized chair-like transition state carrying partial charges at the C-O bond that is broken during the reaction.

Natural CMs belong to two main classes with two distinct folds AroH and AroQ, which are equally efficient, with typical *k*_cat_/*K*_m_ values in the range of 1-5 × 10^5^ M^−1^ s^−1^ (12). The AroH fold, exemplified by the *Bacillus subtilis* CM, has a trimeric pseudo α/β-barrel structure (13,14), whereas the structures of AroQ enzymes have all-α-helical folds (15–21). The AroQ family is further divided into four subfamilies, α-δ (20,21). The AroQ_δ_ subfamily shows abnormally low catalytic activity compared to prototypical CM enzymes. In fact, the first discovered AroQ_δ_ enzyme, the intracellular CM from *Mycobacterium tuberculosis* (MtCM) (20,21), is on its own only a poor catalyst (*k*_cat_/*K*_m_ = 1.8 × 10^3^ M^−1^ s^−1^; (21)), despite its crucial role for producing the aromatic amino acids Tyr and Phe. However, this low activity can be boosted more than a 100-fold to a *k*_cat_/*K*_m_ of 2.4 × 10^5^ M^−1^ s^−1^ through formation of a non-covalent complex with the first enzyme of the shikimate pathway, DAHP synthase (MtDS) (Fig. 1A) (21).

**Figure 1.**
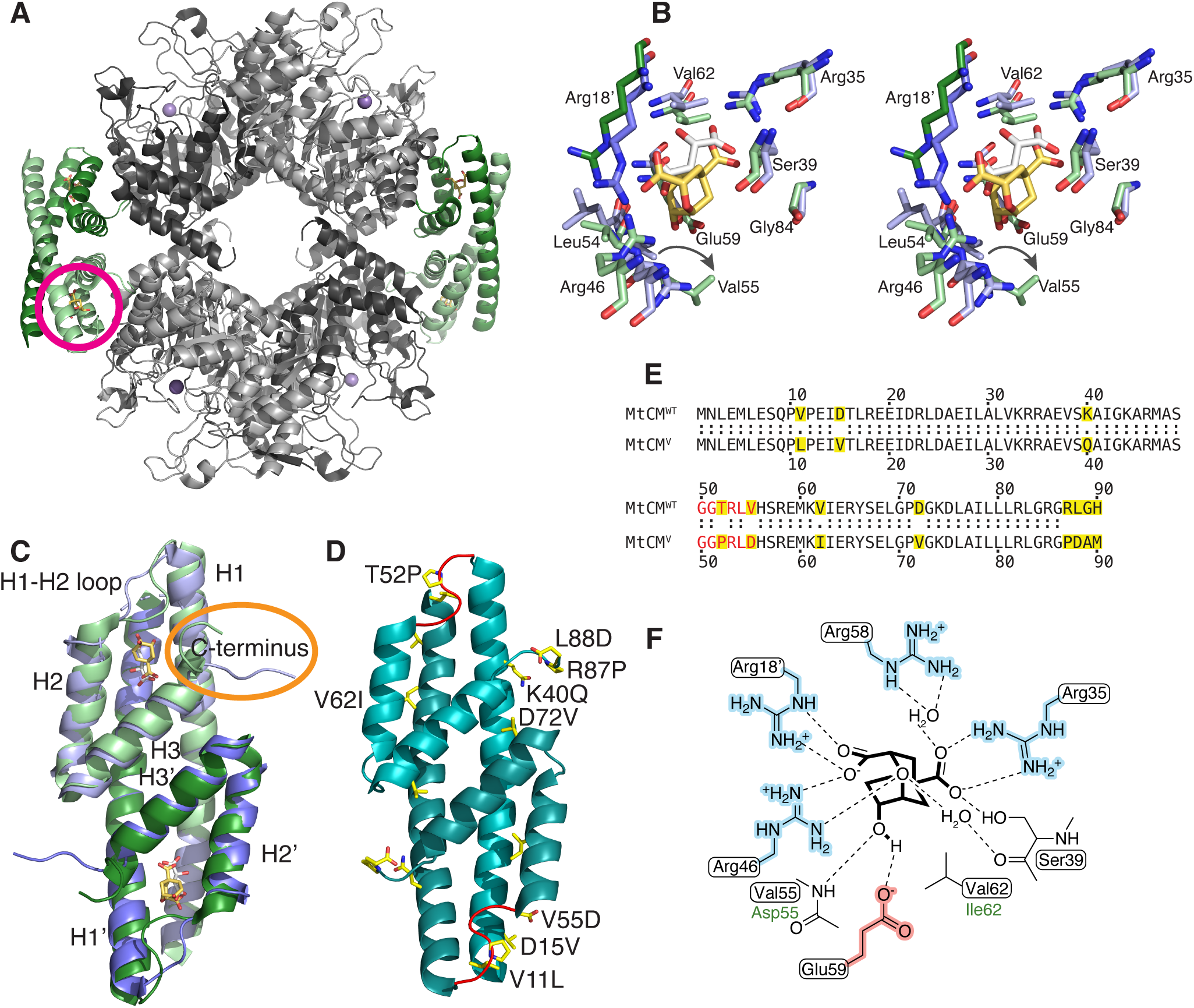
Structural information on *M. tuberculosis* chorismate mutase. (**A**) Cartoon illustration of the heterooctameric complex of MtCM with DAHP synthase (MtDS) (PDB ID: 2W1A (20)). MtCM is colored in shades of green and MtDS in shades of grey to emphasize individual subunits; Bartlett’s transition state analog (TSA) (29) is shown with golden sticks. The location of one of the four active sites of MtCM is marked with a circle. (**B**) Stereo superimposition of CM active sites of MtCM (shades of violet, with malate bound, white sticks) (PDB ID: 2VKL (20)) and of the MtCM-MtDS complex (with TSA bound, golden sticks) (PDB ID: 2W1A (20)), showing several active site residues as sticks. Shades of violet/green (and prime notation for Arg18’) illustrate separate protomers of MtCM/MtCM-MtDS structures, respectively. An arrow shows the shift in position of Val55 upon MtDS binding, allowing H-bond formation of its backbone to TSA. (**C**) Cartoon superimposition of MtCM (PDB ID: 2VKL (20); violet, with white sticks for malate ligand) and activated MtCM (MtCM^DS^) from the MtCM-MtDS complex (PDB ID: 2W1A (20); green, with golden sticks for TSA). The biggest structural changes upon activation are a kink observed in the H1-H2 loop and interaction of the C-terminus (circled in orange) with the active site of MtCM. (**D**) Cartoon representation of the artificially evolved MtDS-independent super-active MtCM variant N-s4.15 (PDB ID: 5MPV (28); cyan), dubbed MtCM^V^ in this work, having a *k*cat/*K*m typical for the most efficient CMs know to date (28). Amino acid replacements accumulated after four cycles of directed evolution are emphasized as yellow side chain sticks (A89 and M90 are not resolved) and labeled for one of the protomers. The H1-H2 loop (shown in red) adopts a kinked conformation similar to that observed for the MtDS-activated MtCM^DS^ shown in (C). (**E**) Sequence alignment of wild-type MtCM (MtCM^WT^) and the highly active variant N-s4.15 (MtCM^V^) (28). Substituted residues are highlighted in yellow, the H1-H2 loop is colored red. (**F**) Schematic representation of the active site of MtCM with bound TSA. Boxed residues refer to the wild-type enzyme, green font color (Asp55, Ile62) to those substituted in MtCM^V^. Charged residues are highlighted in red and blue.

The active site of AroQ CMs is dominated by positive charges, contributed by four arginine residues (Fig. 1F). In MtCM, these are Arg18’, Arg35, Arg46 and Arg58 (with the prime denoting a different MtCM protomer). Of particular importance for catalysis is Arg46 (21), or its corresponding cationic residues in other CMs (of both AroH and AroQ families) (22). However, high catalytic prowess is only achieved when this cationic residue is optimally positioned, such that it can stabilize the developing negative charge at the ether oxygen in the transition state (Scheme 1) (11,14,21,23–25). In MtCM, this is not the case unless MtCM is activated by MtDS (21). The MtDS partner repositions residues of the C-terminus of MtCM for interaction with the H1-H2 loop of MtCM that covers its active site, thereby inducing a characteristic kink in this loop (orange circle in Fig. 1C). This interaction leads to a rearrangement of active site residues to catalytically more favorable conformations (Fig. 1B) (21) and is likely a key contributing factor for the increase in CM activity, as shown by randomizing mutagenesis of the C-terminus followed by selection for functional variants (26). Complex formation also endows MtCM with feedback regulation by Tyr and Phe through binding of these effectors to the MtDS partner (21,27,28). Such inter-enzyme allosteric regulation (28) allows for dynamic adjustment of the CM activity to meet the changing needs of the cell.

The naturally low activity of MtCM in the absence of its MtDS partner enzyme also provided a unique opportunity for laboratory evolution studies aimed at increasing MtCM efficiency. After four major rounds of directed evolution, the highly active MtCM variant N-s4.15 emerged (12), which is abbreviated as MtCM^V^ in this manuscript. This variant showed autonomous CM activity (*k*_cat_/*K*_m_ = 4.7 × 10^5^ M^−1^ s^−1^) twice exceeding that of wild-type MtCM in the MtCM-MtDS complex, and can no longer be activated further through the addition of MtDS (12). The biggest gains in catalytic activity were due to replacements T52P and V55D in the H1-H2 loop and R87P, L88D, G89A, and H90M at the C-terminus (Fig. 1C-E). Of these residues, Pro52 and Asp55 are conserved in the H1-H2 loop of naturally highly active CMs, such as the prototypic CMs from the α- and γ-AroQ subclasses, *i.e.*, EcCM from *Escherichia coli* and *MtCM, the secreted CM from *M. tuberculosis*, respectively (12). The single amino acid exchange that had the largest beneficial effect on activity was V55D (12-fold enhancement of *k*_cat_/*K*_m_), followed by T52P (6-fold gain) (12). Combined, these two mutations, discussed in detail in a previous publication, gave a *k*_cat_/*K*_m_ that was 22 times higher compared to wild-type MtCM (12). The four C-terminal mutations together increased the activity more modestly (by a factor of 4), and the five mutations introduced in the two final evolutionary rounds yielded an additional factor of 5. The resulting combination of large-impact and more subtle residue substitutions in MtCM^V^ (Fig. 1D-E) gave a *k*_cat_/*K*_m_ about 500 times greater than that of the parental starting point (12).

The crystal structure of MtCM^V^ revealed a strongly kinked conformation of the H1-H2 loop. This is reminiscent of the conformation adopted by MtCM when in the complex with MtDS (the crystal structure of MtDS-bound MtCM is in the following referred to as MtCM^DS^), and differs considerably from that observed in free MtCM (Fig. 1C-D) (12). However, in the crystal structures of free wild-type and top-evolved MtCM^V^, both the H1-H2 loop and the C-terminus are involved in extensive crystal contacts, making an unbiased structural evaluation of the mutational effects in these parts of the enzyme impossible. In solution, these regions are assumed to be more flexible compared to the α-helical segments of MtCM.

Here, we used molecular dynamics (MD) simulations to investigate the behavior of MtCM in the absence or presence of ligands, and to analyze whether the protein is able to interconvert between activated and non-activated conformations in the absence of the MtDS partner enzyme. We also compared the wild-type MtCM with the evolved MtCM^V^, to see if the acquired amino acid substitutions introduced any new interactions or if they altered the probabilities of existing ones, with potential impact on catalytic activity. From an assessment of the dynamic properties of MtCM and MtCM^V^, we proposed a set of single, double and triple C-terminal variants of the enzyme and subsequently tested these experimentally.

## MATERIALS AND METHODS

### Construction of Untagged MtCM Variants

General cloning was carried out in *Escherichia coli* DH5α or XL1-Blue (both Stratagene, La Jolla, CA, USA). All cloning techniques and the bacterial culturing were performed according to standard procedures (29). Oligonucleotide synthesis and DNA sequencing were performed by Microsynth AG (Balgach, Switzerland).

For construction of expression plasmids pKTCMM-H-V55D and pKTCMM-H-T52P for the native MtCM single variants, the individual site-directed mutants were first constructed in the pKTNTET background (providing an N-terminal His_6_ tag, first 5 residues missing). Parts of the MtCM gene were amplified using oligonucleotides 412-MtCM-N-V55D (5’- GTTCGCTAGCGGAGGTACACGTTTGGATCATAGTCGGGAGATGAAGGTCATCGAAC) or 413-MtCM-N-T52P (5’- GTTCGCTAGCGGAGGTCCGCGTTTGGTCCATAGTCGGGAGATGAAGGTCATCGAAC) together with oligonucleotides 386-LpLib-N2 (5’- GGTTAAAGCTTCCGCAGCCACTAGTTATTAGTGACCGAGGCGGCCACGGCCCAAT) on template pMG248 (12) to create a 163 bp PCR product. The PCR products were restriction digested with *Nhe*I and *Hin*dIII and the resulting 148 bp fragments were individually ligated to the accordingly cut 2873 bp fragment from acceptor vector pKTNTET-0 (12). The ligation was performed with T4 DNA ligase (New England Biolabs, Ipswich, MA, USA) overnight at 16°C. The ligation products were transformed into chemically competent *E. coli* XL1-Blue cells. The cloned PCR’ed DNA fragments were confirmed by Sanger sequencing. Subsequently, the genes for MtCM-T52P and MtCM-V55D were isolated by restriction digestion using enzymes *Xho*I and *Spe*I followed by a preparative agarose gel, yielding corresponding 260 bp fragments. pKTCMM-H (21) was used as acceptor vector and was accordingly restriction digested with *Xho*I and *Spe*I, yielding a 4547 bp acceptor fragment. The fragments were ligated overnight at 16°C, using T4 DNA ligase. The ligation products were transformed into chemically competent *E. coli* KA12 cells and the inserts were analyzed by Sanger sequencing. The gene for variant PHS10-3p3 (12), carrying an N-terminal His_6_-tag and missing the first 5 residues, was recloned into the native format provided by plasmid pKTCMM-H. Acceptor vector pKTCMM-H and pKTNTET-PHS10-3p3 were restriction digested with *Xho*I and *Spe*I and the fragments isolated from preparative agarose gels. The 4547 bp and 260 bp fragments were ligated overnight at 16°C with T4 DNA ligase and transformed into chemically competent XL1-Blue cells. The relevant gene sequence was confirmed by Sanger sequencing.

Different C-terminal variants of the MtCM gene were generated by PCR mutagenesis. DNA fragments were amplified with the same forward primer (containing an *Nde*I site, underlined) and different reverse primers (containing an *Spe*I site, underlined) on different DNA templates. The gene encoding MtCM L88D was produced by PCR with primers LB5 (5’-TCCGCA<u>CATATG</u>AACCTGGAAATG) and LB4 (5’-TAAGCA<u>ACTAGT</u>TATTAGTGACCGTCGCG) on the template plasmid pKTCMM-H carrying the wild-type gene (21). The gene for the triple variant MtCM (T52P V55D L88D) was assembled with primers LB5 and LB4 on a pKTCMM-H derivative containing MtCM variant 3p3 (T52P V55D) (12). The gene for MtCM variant PNAM (D88N) was generated with primers LB5 and LB6 (5’- TAAGCA<u>ACTAGT</u>TATTACATAGCATTCGGA), and for the MtCM variant PLAM (D88L) with primers LB5 and LB7 (5’- TAAGCA<u>ACTAGT</u>TATTAGTGACCAAGCGGA), in both cases using a version of the template plasmid pKTCMM-H, into which the gene for the top-evolved s4.15 variant had been inserted (12). The resulting 296 bp PCR fragments containing *Nde*I and *Spe*I restriction sites at the 5’ and 3’ ends of the MtCM gene, respectively, were digested with the corresponding enzymes to yield 279 bp fragments. These fragments were ligated to the 4528 bp *Nde*I-*Spe*I fragment of pKTCMM-H yielding the final 4807 bp plasmids.

### Protein Production and Purification

*E. coli* strain KA13 (18,30) carrying an endogenous UV5 P*_lac_*-expressed T7 RNA polymerase gene was used to overproduce the (untagged) MtCM variants. KA13 cells were transformed by electroporation with the appropriate pKTCMM-H plasmid derivative that carries the desired MtCM gene variant.

For the two crystallized MtCM variants T52P (MtCM^T52P^) and V55D (MtCM^T52P^) the transformed cells were grown in baffled flasks at 30 °C in LB medium containing 100 µg/mL sodium ampicillin until the OD_600_ reached 0.5. Gene expression was induced through the addition of isopropyl-β-D-thiogalactopyranoside (IPTG) to a final concentration of 0.5 mM, and incubation was continued overnight. The cells were harvested by centrifugation (6500 g for 20 min at 4 °C) and frozen at −80 °C before being resuspended in a buffer suitable for ion exchange chromatography, supplemented with DNase I (Sigma), 150 µM phenylmethanesulfonyl fluoride (PMSF) and cOmplete protease inhibitor cocktail (Roche). Cells were lysed using BeadBeater (BioSpec BSP 74340, Techtum Lab AB), with four times 30 s pulses with a 60 s wait between each pulse. Insoluble debris was removed by centrifugation (48,000 g for 30 min at 4 °C).

The resuspension buffer was selected based on the theoretical isoelectric point (pI) of the protein. MtCM T52P has a pI of 8.14, so the pellet was resuspended in 50 mM 2-(N-morpholino)ethanesulfonic acid (MES), pH 6.5. MtCM V55D has a pI of 6.74; therefore, the pellet was resuspended in 50 mM acetic acid, pH 5.25. After lysis and centrifugation, the soluble lysate was loaded onto a HiTrap XL SP column (GE Healthcare) for cation exchange chromatography and eluted with a 0-0.5 M NaCl gradient. Purity of the eluted fractions was gauged by SDS PAGE analysis and sufficiently pure fractions were pooled and concentrated using concentrator tubes with a 5 kDa molecular mass cut off (Vivaspin MWCO 5K). The proteins were then further purified by size-exclusion chromatography using a Superdex 75 300/10 column (GE Healthcare) with running buffer 20 mM 1,3-bis[tris(hydroxymethyl)methylamino]propane (BTP), pH 7.5, 150 mM NaCl. Finally, the proteins were concentrated (Vivaspin MWCO 5K), frozen and stored at −80°C.

For the sets of MtCM variants probed for the catalytic impact of particular C-terminal mutations, 500 mL LB medium cultures containing 150 μg/mL sodium ampicillin were inoculated with 5 mL overnight culture of the desired transformant and grown at 37 °C and 220 rpm shaking to an OD_600 nm_ of 0.3 - 0.5. Protein production was induced by addition of IPTG to 0.5 mM and culture growth was continued overnight at 30°C.

The cells were harvested by centrifugation (17,000 g for 10 min at 4 °C) and washed once with 100 mM Tris-HCl, pH 7.5. The cells were pelleted again and the cell pellet either frozen for storage at −20 °C or directly resuspended in 80 mL sonication buffer (50 mM sodium phosphate, 0.3 M NaCl, pH 7.0). Cells were disrupted by sonication on ice (15 min total pulse time with 45 s pulse / 30 s pause cycles at 50% amplitude; Q700 sonicator, QSonica). The crude lysate was cleared by centrifugation (20,000 g for 20 min at 4 °C). The supernatant was supplemented with sonication buffer to 100 mL, 42 g of ammonium sulfate was added, and the solution was stirred at 4 °C for 1.5 h. The precipitate was pelleted by centrifugation (10,000 g for 30 min at 4 °C), dissolved in 8 mL low-salt buffer (20 mM piperazine, pH 9.0), and dialyzed against 1 L low-salt buffer overnight. Dialysis was repeated against another 1 L of low-salt buffer for 3 h before application to a MonoQ (MonoQ HR 10/10, Pharmacia) FPLC column (Biologic Duoflow system, Bio-Rad). The sample was eluted over 80 mL in 20 mM piperazine by applying a gradient from 0% to 30% of a high-salt buffer (20 mM piperazine, 1 M NaCl, pH 9.0).

The MonoQ fractions containing the protein of interest were pooled and concentrated to less than 1 mL. The concentrated sample was directly applied to a gel-filtration column (Superdex Increase 75 10/300 GL, GE Healthcare) and eluted in 20 mM BTP, 150 mM NaCl, pH 7.5. Protein identity was confirmed by LC-MS (MoBiAS facility, Laboratory of Organic Chemistry, ETH Zurich), with the observed mass being withing 1 Da of the calculated mass. Protein purity was assessed by SDS-polyacrylamide gel electrophoresis (PhastGel Homogenous 20 pre-cast gels, GE Healthcare) and the enzyme concentration ([E]) was determined by using the Bradford assay (31).

### X-ray Crystallography

MtCM variants T52P (MtCM^T52P^) and V55D (MtCM^T52P^) were crystallized in 96-well 2-drop MRC crystallization plates (SWISSCI) by the sitting drop vapor diffusion technique. Diffraction-quality crystals of MtCM^T52P^ grew at 20 °C from a 1:1 (375 nL + 375 nL) mixture of protein (28 mg/mL in 20 mM Bis Tris propane, pH 7.5) and reservoir solution containing 0.2 M sodium malonate, 20% PEG 3350 (w/v) and 0.1 M Bis Tris propane buffer, pH 8.5 (PACT *premier* crystallization screen, condition H12; Molecular Dimensions Ltd.). Crystals of MtCM^V55D^ were obtained from a 1:1 (375 nL + 375 nL) mixture of protein (44 mg/mL in 20 mM Bris Tris propane, pH 7.5, 150 mM NaCl) and reservoir solution containing 0.2 M zinc acetate dihydrate, 10% w/v PEG 3000, and 0.1 M sodium acetate, pH 4.5, (JCSG-plus crystallization screen, condition C7; Molecular Dimensions Ltd.) at 20 °C.

Diffraction data of MtCM^T52P^ and MtCM^V55D^ crystals were collected at the European Synchrotron Radiation Facility (ESRF, Grenoble, France), at the ID30A-3/MASSIF-3 (Dectris Eiger X 4M detector) and ID29 (Pilatus detector) beamlines, respectively, covering 120° with 0.1° oscillation. Diffraction images were integrated and scaled using the *XDS* software package (32); merging and truncation were performed with *AIMLESS* (33) from the *CCP4* program suite (33,34). Since data collection statistics of both crystals suggested the presence of anisotropy, the *XDS* output was reprocessed for anisotropy correction and truncation using the *STARANISO* server (35). The ‘aniso-merged’ output files (merged MTZ file with an anisotropic diffraction cut-off) were subsequently used for structure solution and refinement (Table 1).

**Table 1:** RMSF values of selected active site residues

|  | RMSF (Å) |  |  |  |
| --- | --- | --- | --- | --- |
|  | MtCM<sup>WT</sup> | MtCM<sup>V</sup> | MtCM<sup>LC</sup> | MtCM<sup>V55D</sup> |
| Arg18' | 2.2 ± 0.5 | 1.5 ± 1.0 | 0.7 ± 0.2 | 1.1 ± 0.5 |
| Arg35 | 0.8 ± 0.4 | 0.7 ± 0.4 | 0.4 ± 0.1 | 0.6 ± 0.2 |
| Arg46 | 2.0 ± 0.6 | 1.7 ± 0.7 | 0.7 ± 0.5 | 0.9 ± 0.5 |
| Arg58 | 2.3 ± 1.0 | 1.9 ± 2.0 | 2.0 ± 0.9 | 1.5 ± 0.6 |

RMSF values were calculated as an average over all non-hydrogen atoms for each residue compared to the average structure of the simulation.

The crystal structures of MtCM^T52P^ and MtCM^V55D^ were solved by molecular replacement with the program *PHASER* MR (36). The structure of the top-evolved MtCM variant MtCM^V^ (PDB ID: 5MPV (12)) was used as a search model for solving the structure of MtCM^T52P^ since it was expected to be a better match at the Pro52-containing H1-H2 loop compared to wild-type MtCM. For MtCM^V55D^, we used the MtCM structure from the MtCM-MtDS complex (PDB ID: 2W1A (21)) as a search model, after truncation of the termini and the H1-H2 loop, and removal of the ligand.

The two structures were subsequently refined, alternating between real-space refinement cycles using *Coot* (37) and maximum likelihood refinement with *REFMAC5* (38). The models were improved step-wise by first removing ill-defined side chains, and subsequently adding missing structural elements as the quality of the electron density map improved. Water molecules and alternative side chain conformations were added to the MtCM^T52P^ model towards the end of the refinement process, where positive peaks in the σ_A_-weighted *F*o-*F*c difference map and the chemical surroundings allowed for their unambiguous identification. As a last step, occupancy refinement was carried out with phenix.refine, a tool of the *PHENIX* software suite (39). The final structure of MtCM^T52P^ was deposited in the Protein Data Bank (PDB) (40) with deposition code 6YGT. Data collection and refinement statistics are summarized in Supporting Table S1.

### Determination of Enzyme Kinetic Parameters

Michaelis-Menten kinetics of the untagged purified MtCM variants were determined by a continuous spectroscopic chorismate depletion assay (Lambda 20 UV/VIS spectrophotometer, Perkin Elmer). The enzymes were diluted to working stock solutions in 20 mM potassium phosphate, pH 7.5, containing 0.01 mg/mL bovine serum albumin. The assays were performed at 30 °C in either 50 mM potassium phosphate, pH 7.5, or 50 mM BTP, pH 7.5. Different chorismate concentrations ([S]) ranging from 10 to 1500 μM were used at 274 nm (ε_274_ = 2630 M^−1^ cm^−1^) or 310 nm (ε_310_ = 370 M^−1^ cm^−1^). Chorismate disappearance upon enzyme addition was monitored to determine the initial reaction velocity (*v*_0_). The obtained data was fitted to the Michaelis-Menten equation (with the program KaleidaGraph (Synergy Software, Reading, PA, USA) to obtain the catalytic parameters *k*_cat_ and *K*_m_.

### Molecular Dynamics Simulations

Molecular dynamics (MD) simulations were carried out on a number of representative structures for CM. They included two independent sets of simulations for apo MtCM, starting either from the X-ray crystal structure of MtCM in complex with malate (after removing malate) (PDB ID: 2VKL (21)), or from the structure of the CM polypeptide in the apo MtCM-MtDS complex (PDB ID: 2W19 (21), chain D). The malate complex was chosen over ligand-free MtCM (PDB ID: 2QBV (41)) due to its higher resolution and better refinement statistics. Both simulations gave essentially the same result; therefore, we will not refer to the second data set any further. For the highly active evolved MtCM variant (MtCM^V^), we used the recent crystal structure (PDB ID: 5MPV (12)). The MtCM-ligand complex (MtCM^LC^) was taken from PDB ID: 2W1A (21), excluding the MtDS partner protein, where MtCM was co-crystallized with a transition state analog (TSA) in its active site. Finally, the V55D variant was modeled based on a partially refined experimental structure (Table S1). Residues that were not fully defined were added to the models using (often weak) electron density maps as reference in *Coot* (37). When no interpretable density was visible, geometric restraints (and α-helical restraints for residues in helix H1) were applied during model building, to ensure stable starting geometries. The N-termini of all the models were set at Glu13, corresponding to the first defined residue in almost all the resolved structures available. Glu13 was capped with an acetyl group to imply the continuation of the H1 helix. CM dimers were generated by 2-fold crystallographic symmetry.

Missing H-atoms were added to the model and the systems were solvated in a periodic box filled with explicit water molecules, retaining neighboring crystallographic waters, and keeping the protein at least 12 Å from the box boundaries. The systems were neutralized through the addition of Cl^−^ ions at a minimum distance of 7 Å from the protein and each other. Additional buffering moieties like glycerol or sulfate ions found in the crystals were not considered. MD simulations were run using the Gromacs 5.1.4 package (42,43) using the AMBER 12 force fields (44,45) for the protein moieties (44,45) and the TIP3P model for water (46). The ligand was modeled using the GAFF force field (47). The smooth Particle Mesh Ewald method was used to compute long-range electrostatic interactions (48), while a cut-off of 11 Å was used to take in to treat the Lennard-Jones potential.

The systems were minimized using the steepest descent/conjugate gradients algorithms for 500/1500 steps until the maximum force was less than 1000 kJ mol^−1^ nm^−1^. To equilibrate and heat the systems, first we ran 100 ps MD in the NVT ensemble starting from a temperature of 10 K, using the canonical velocity rescaling thermostat (49) followed by 100 ps in the NpT ensemble with a Parrinello-Rahman barostat (50) targeting a final temperature of 310 K and a pressure of 1 atm. After initial equilibration, 1 µs of MD simulations were performed for each system. In all MD simulations the time step size was set to 2 fs.

## RESULTS

The fact that MtCM exhibits only low natural catalytic activity provided us with a perfect opportunity to probe features that optimize CM catalysis by directed evolution (12). Since the biggest gains in catalytic activity were contributed by exchanging the H1-H2 loop residues 52 (T52P) and 55 (V55D), we set out to determine the crystal structures of these two enzyme variants. These two substitutions led to a combined increase in *k*_cat_/*K*_m_ by 22-fold compared to the parent enzyme (12).

### Crystal Structures of MtCM^T52P^ and MtCM^V55D^

Whereas MtCM^T52P^ crystals had the same space group (*P*4_3_2_1_2) and similar cell parameters as the wild-type enzyme (PDB IDs: 2VKL(21) and 2QBV(41)), with one protomer in the asymmetric unit, MtCM^V55D^ crystallized in a different space group (*P*22_1_2_1_), where the asymmetric unit contained the biological dimer. The MtCM^T52P^ structure was refined to 1.6 Å and *R*_work_/*R*_free_ values of 24.0/26.5% (Table S1; Fig. S1B), whereas MtCM^V55D^ diffraction data yielded lower-quality electron density, particularly at the H1-H2 loop (Fig. S1C and D). Consistent with this, the Wilson *B*-factor of MtCM^V55D^ is high (57.8 Å^2^), indicating structural disorder. Refinement of the MtCM^V55D^ model stalled at *R*_work_/*R*_free_ values of 27.6/34.9%, with very high *B*-factors for active site H1-H2 loop residues, especially for protomer B. For both structures, residues preceding residue Glu13 and C-terminal to Leu88 showed poorly defined electron density. Therefore, the terminal residues were not included in the final model.

Overall, the crystal structures of both MtCM^T52P^ (PDB ID: 6YGT) and MtCM^V55D^ are very similar to the structure of substrate-free wild-type MtCM (PDB ID: 2QBV (41)), with RMSD = 0.3 Å and 0.4 Å, respectively. However, the active site H1-H2 loops (^47^MASGGPRLDHS^57^) of both protomers of MtCM^V55D^ adopt a different conformation (RMSD = 2.3 Å compared to PDB ID: 2QBV), which most closely resembles the kinked conformation in the MtCM-MtDS complex (PDB ID: 2W19 (21); RMSD = 0.8 Å) (Fig. 2A). In the crystal structure of MtCM^V55D^, Asp55 in the H1-H2 loop forms a salt bridge with Arg46, similar to the one in MtCM^V^ (compare Figs. S1G-H with S1E). This interaction pre-organizes the active sites of both MtCM variants for catalytic activity, mimicking MtCM in the complex with MtDS (Fig. 2B). However, the overall conformation of the active site loop, which is involved in extensive crystal contacts that are highly distinct for the different crystal forms (Fig. S2), differs significantly between the structures (Figs. S1 and 2A).

**Figure 2.**
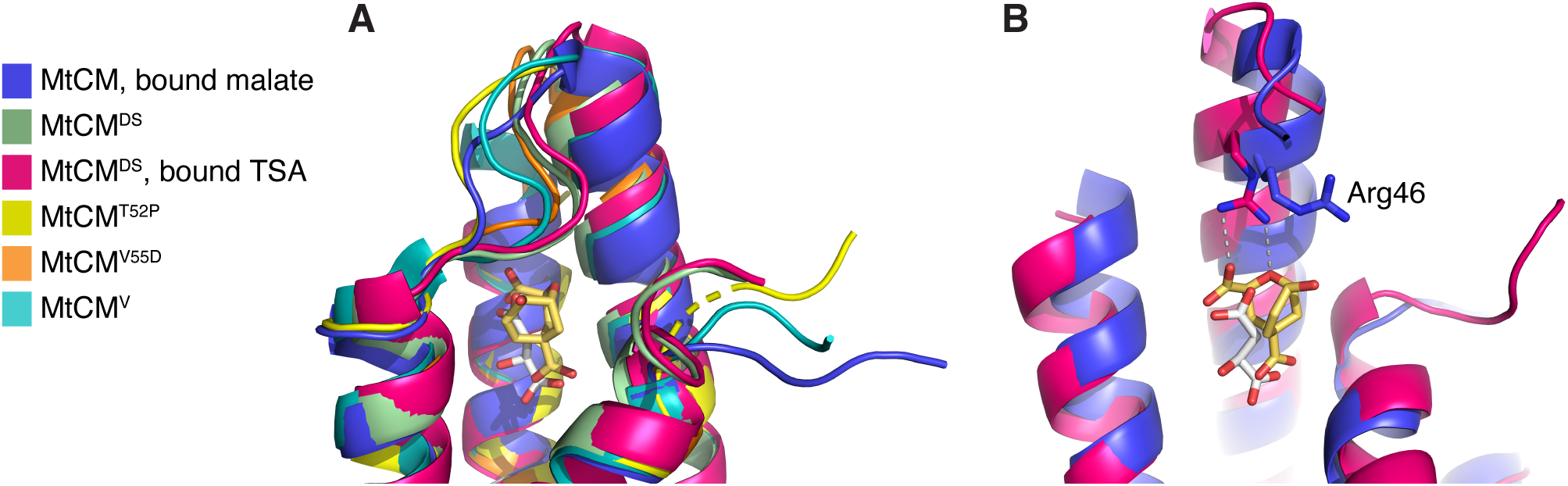
Comparison of MtCM crystal structures, with focus on Arg46 and H1-H2 loop. (**A**) Superimposition of the active site of MtCM (PDB ID: 2VKL (20); violet), MtCM^DS^ (PDB ID: 2W19 (20); green), MtCM^DS^-TSA complex (PDB ID: 2W1A (20); pink), MtCM^T52P^ (PDB ID: 6YGT, this work; yellow), MtCM^V55D^ (this work; orange), and top-evolved MtCM^V^ (PDB ID: 5MPV (28); cyan). Shown is the cartoon representation featuring the H1-H2 loop, with the ligands depicted as sticks. (**B**) Superimposition of MtCM (PDB ID: 2VKL (20); violet, with bound malate in grey sticks) and MtCM in the MtCM-MtDS complex (PDB ID: 2W1A (20); pink, with TSA in golden sticks, corresponding to MtCM^LC^) in cartoon representation, with the catalytically important Arg46 depicted as sticks. MtDS binding promotes the catalytically competent conformation of Arg46. Helix H2 was removed for clarity.

### MD Simulations

To evaluate the behavior of MtCM in the absence of crystal contacts, we probed the MtCM structures by MD simulations. We used four model systems: low-activity apo wild-type MtCM, MtCM^LC^ (‘Ligand Complex’: wild-type MtCM from the MtCM-MtDS structure in complex with TSA, the transition state analog of the CM reaction (51); Scheme 1 and Fig. 1), MtCM^V^, corresponding to the highly active evolved variant N-s4.15 (12), and MTCM^V55D^, which shows the highest catalytic activity among the single-substitution MtCM variants (12). We compared the overall dynamic profiles of these models and inspected the interactions formed between the C-termini and the H1-H2 loops covering the active sites, to find general features that could be associated with increased catalytic competence.

#### Apo structures of MtCM are characterized by significant flexibility

We anticipated that the model systems would more or less retain the same fold as observed in the crystal structures, but that regions associated with crystal contacts, like the C-termini and the H1-H2 loop, would rapidly move away from their starting positions. Instead, the MD simulations revealed large changes from the initial crystal geometries in the apo protein structures, causing a rather high root-mean-square-deviation (RMSD) from the original crystal structure geometry for the CM core regions (RMSD = 2.8 ± 1.2 Å (MtCM) or 3.4 ± 1.5 Å (MtCM^V^)). In particular, helix H2 showed a tendency to unravel (Fig. 3). Due to the large flexibility observed, the two protomers making up the biological dimer instantaneously broke their symmetry, independently exploring different conformations in two chains. In contrast, the ligand-bound structure MtCM^LC^ retained the secondary structure throughout the 1-µs simulation (Fig. 3), with a lower RMSD (1.7 ± 0.6 Å) than the two apo structures. Intriguingly, a similar stabilization was observed for the unliganded variant MtCM^V55D^ (Fig. 3).

**Figure 3.**
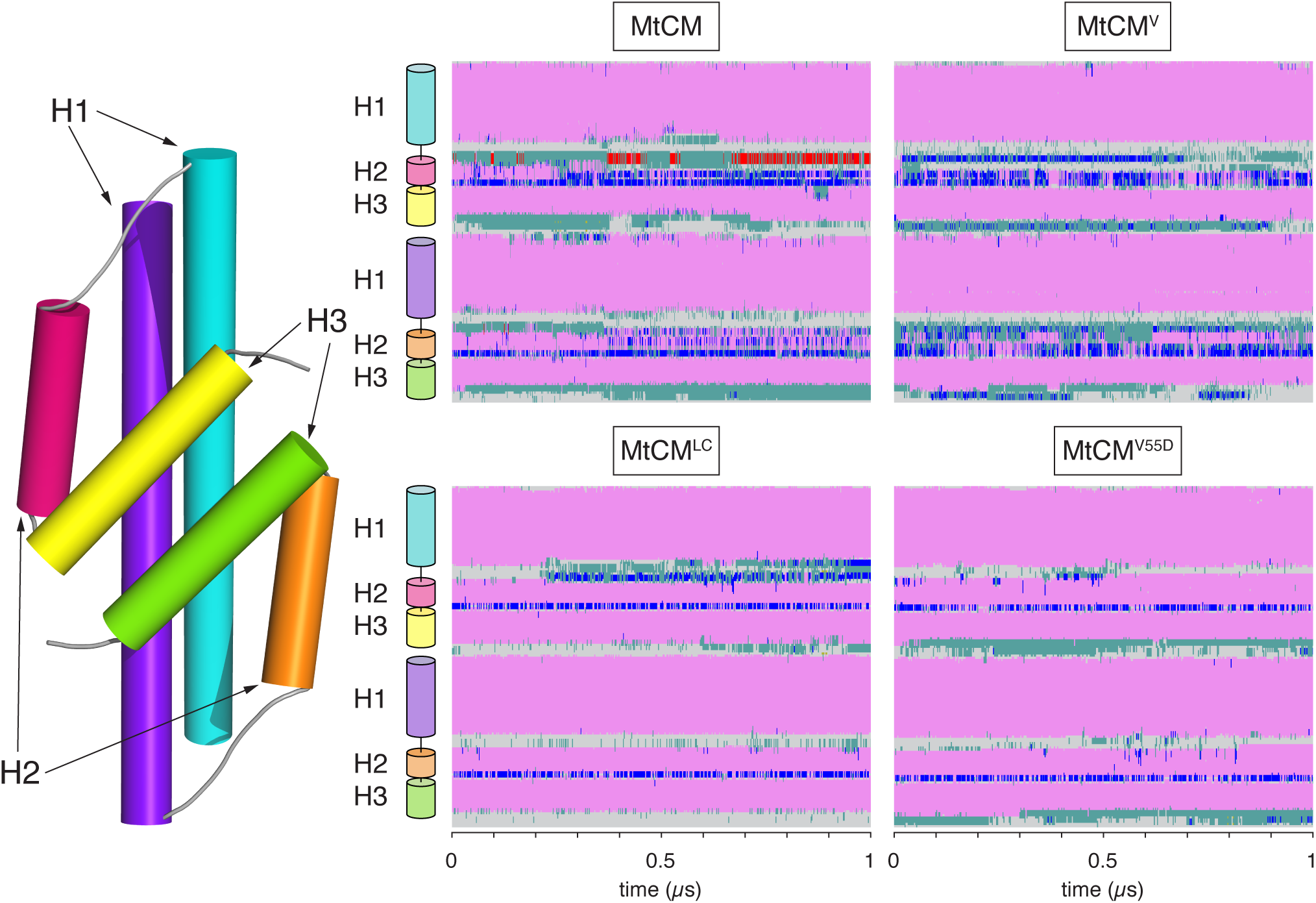
Secondary structure changes of MtCM during MD simulations. Estimated secondary structure of MtCM over 1 µs of MD simulation. Color code is: α-helical structure (magenta), 310 helix (blue), π-helix (red), turn (green), coil (gray). The top panels report data for apo MtCM and MtCM^V^ systems, showing clear instability of H2. The bottom panels present the data for the holo-MtCM^LC^ system and for the single V55D variant (apo structure), which in contrast retained all secondary structure elements within the simulation time.

#### Kinked conformation of the H1-H2 loop

One of the biggest conformational changes in the crystal structure upon formation of the MtCM-MtDS complex occurs in the H1-H2 loop (Fig. 1C and 2A) (21). Whereas in the X-ray structure of the MtDS-activated MtCM, the H1-H2 loop is strongly kinked, this is not the case in non-activated MtCM. We investigated the conformational landscape of this loop by simulations, using Arg53 from the loop as reporter residue. As shown in Figure 4, in one of the two protomers of MtCM, Arg53 remained in an extended conformation for the entire 1-µs MD simulation. In contrast, the same amino acid in the other protomer oscillated between the extended and the helical region of the Ramachandran plot (Fig. 4), the latter being characteristic of the catalytically active conformation of the loop. Statistically averaging the two distributions, it appears that the apo form of MtCM is preferentially found in its inactive conformation, whereas in MtCM^V^ both protomers assumed the kinked active loop conformation, and retained it for the whole length of the simulation. However, TSA binding promoted the active conformation also in wild-type MtCM (represented by MtCM^LC^). The fact that the fluctuations of the MtCM^V^ H1-H2 loop are contained within the conformational basin of the catalytically competent geometry (Fig. 4, Table 1) is an indication that MtCM^V^ has an intrinsically pre-organized loop, a condition that helps to minimize the entropy loss during substrate binding and consequently favors catalysis.

**Figure 4.**
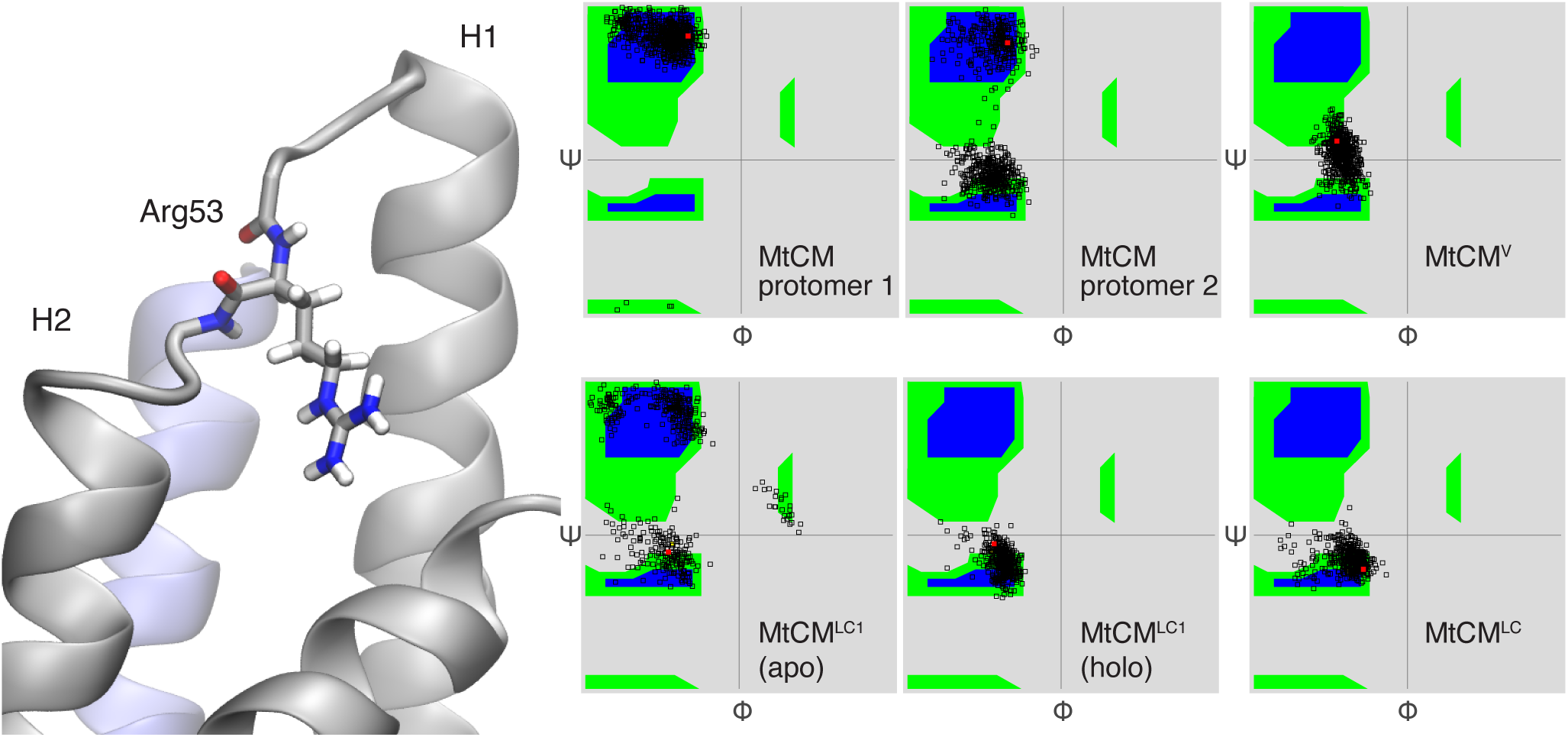
Conformation of Arg53 in the H1-H2 loop. Ramachandran plot showing backbone dihedral angles Φ and ψ for Arg53. Red dots mark starting conformations. For MtCM, the H1-H2 loops from the two protomers (left and middle plots in the top row) assume different ensembles of conformations; overall, the catalytically favored conformation (Ψ ∼ 0) is observed less frequently than the nonproductive one. In contrast, in MtCM^V^ both loops retained the active conformation during the entire course of the simulation, similarly to what was observed for MtCM^LC^. When only one ligand bound (MtCM^LC1^), the TSA-loaded site (holo) retained the active conformation, while the loop in the other protomer (apo) remained flexible.

To test the effect of ligand binding, we repeated simulations of MtCM loaded with only one TSA ligand (MtCM^LC1^). Interestingly, ligand presence in one of the two binding pockets was sufficient to stabilize the structure of the whole dimer. Nonetheless, the H1-H2 active site loop of the apo protomer retained its intrinsic flexibility (Fig. 4). The fact that the active site loop of the unloaded protomer behaved like the apo MtCM system suggests that the two active sites in MtCM retain considerable independence.

Contrary to MtCM and MtCM^V^ (Fig. 1C/D), the presence of the additional carboxylate group in MtCM^V55D^ promoted the elongation of helix H2, resulting in a significant shortening of the active H1-H2 loop (Fig. 5A). This structural rearrangement is associated with the formation of persistent salt bridges between Asp55, now localized in the first turn of H2, and active site residues Arg18’ and Arg46 that are retained for the entire length of the simulation.

**Figure 5.**
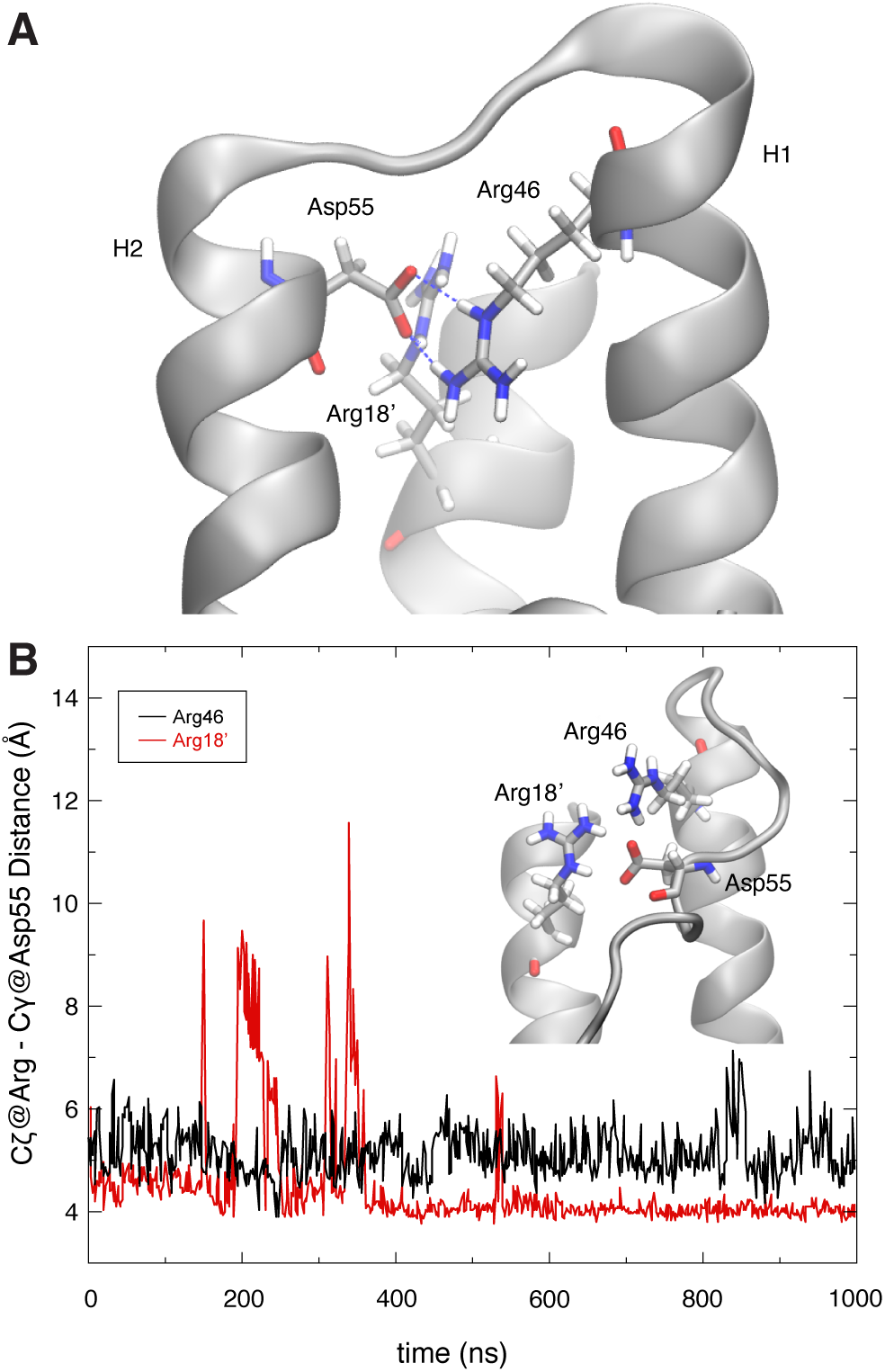
Role of MtCM^V^ residue Asp55 in positioning active site residues. (**A**) Extension of H2 and stabilization of the H1-H2 loop by residue Asp55. Substitution of Val55 by Asp stabilizes helix H2 through interactions with Arg18’ and Arg46 across the active site (the image shows the structure of V55D after 1 μs of MD simulations). Note that Arg46 is a catalytically essential residue for MtCM and its correct orientation critical for catalytic proficiency. (**B**) Shown is the distance plotted between MtCM^V^ Arg46 (black, chain A) or Arg18’ (red, from chain B) and Asp55 (chain A) observed during the simulation. In both cases, the distance measured is between Asp Cγ and Arg C*ζ*, using PDB nomenclature.

#### Positioning of active site residues

The MtCM active site contains four arginine residues (Fig. 1F), among them the key catalytic residue Arg46. In contrast to the observation in the two MtCM-MtDS crystal structures, the conformation of Arg46 was not strictly maintained during MD simulations. In the absence of a ligand, Arg18’, Arg46, and Arg58 repelled each other, and at least one of the residues was pushed out of the active site in the majority of the simulations. Only one of the four arginine residues (Arg35) maintained its position (Table 1), appropriatedly placed for substrate binding by wild-type MtCM. This changes upon complex formation with MtDS, guiding also the important Arg46 into a catalytically competent conformation.

In contrast to MtCM^WT^, the two variants MtCM^V^ and MtCM^V55D^ exhibited lower RMSF values for all active site Arg residues (Table 1; Fig. 1F) and maintained their catalytically competent conformation during the MD simulations even in the apo forms (Fig. 5). The more stable positioning of Arg18’ and Arg46 appears to be a direct consequence of the replacement of Val55 with Asp, which introduces a negative charge, mitigating the surplus positive charges in the active site.

#### Interactions between C-terminal residues and H1-H2 loop

A crucial factor for the enhanced activity of MtCM in the MtCM-MtDS complex is an MtDS-induced interaction between MtCM’s H1-H2 loop and its C-terminus (21). The interaction can be divided into two contributions: A salt-bridge between the C-terminal carboxylate and the side chain of Arg53, and a hydrophobic contact between Leu54 and Leu88 (Fig. S3A).

Our 1-μs long simulations detected persistent, multiple interactions involving the C-terminal carboxylate. In contrast, the hydrophobic contacts between Leu54 and Leu88 were disrupted in the first ns, and almost never observed again during the rest of the simulation time (Fig. S3B/C).

##### Salt bridges with C-terminus

In our MD simulations, the C-terminal carboxylate formed interchangeable contacts with Arg53 and the catalytically important Arg46 (21), which is located in the last turn of helix H1 (Fig. 6A). Notably, the presence of a salt bridge between Arg46 and the C-terminus correlated with the apparently active conformation of the H1-H2 loop (Fig. 6A).

**Figure 6.**
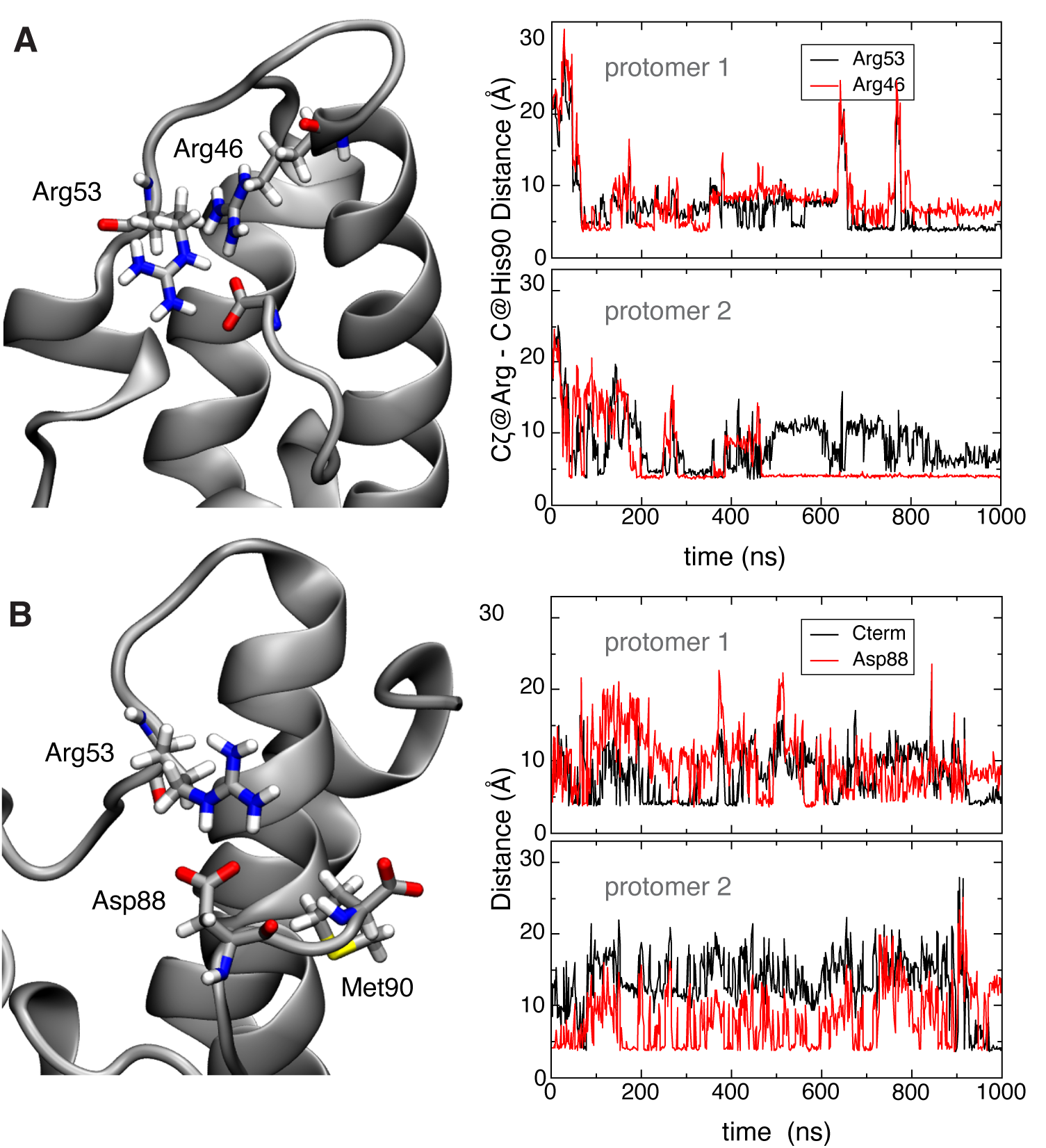
Interaction between C-terminal carboxylate of MtCM and H1-H2 loop. (**A**) The right panels show the distance between the Cζ carbon of arginine residues 53 or 46 and the carboxylate carbon of the C-terminus of MtCM in the two protomers (upper and lower panels). Formation of a steady contact (< 5 Å) with Arg46 (bottom panel) corresponds to stabilization of the catalytically productive H1-H2 loop conformation, which allows for stabilization of the transition state of the chorismate to prephenate rearrangement (Fig. 1B and F). (**B**) Interaction between C-terminal residues and the H1-H2 loop in MtCM^V^. Salt-bridge contacts between Arg53 and the carboxyl groups of Asp88 (red line) and Met90 (C-terminus; black line) in MtCM^V^ in the two protomers. The top and bottom panels on the right show the evolution of the distances over time between Arg53’s Cζ and the corresponding carboxylate carbons for each of the two protomers of MtCM^V^.

The observed fluctuation suggests that the catalytically competent conformation of the binding site is malleable in wild-type MtCM, and that additional interactions, *i.e.*, with the substrate, are required to stabilize it. This is in keeping with studies of a topologically redesigned monomeric CM from *Methanococcus jannaschii*. This artificial enzyme was found to be catalytically active in the presence of the substrate despite showing extensive structural disorder reminiscent of a molten globule (52).

##### MtCM^V^ exhibits strengthened interactions between C-terminus and H1-H2 loop

In MtCM^V^, the four C-terminal residues Arg-Leu-Gly-His (RLGH) are substituted with Pro-Asp-Ala-Met (PDAM) at positions 87-90, which include another carboxylate, introduced through Asp88. Our MD simulations show that the Asp88 carboxylate in the evolved variant MtCM^V^ offers an alternative mode of interaction with Arg53 of the H1-H2 loop (Fig. 6B), which is not possible for wild-type MtCM (Fig. S4). This allows for a persistent interaction of C-terminal residues with the H1-H2 loop throughout the simulation, while maintaining a highly flexible C-terminus. Moreover, in MtCM^V^ Arg46 is topologically displaced from its original position with respect to the loop, and no longer able to engage in a catalytically unproductive salt bridge with the C-terminus.

Another interesting substitution, which emerged within the four C-terminal residues during the laboratory evolution towards variant MtCM^V^, was a proline residue (RLGH to PDAM) (12). However, in contrast to Pro52, Pro87 did not appear to have a major influence on the simulations. While Pro52 is likely contributing to H1-H2 loop rigidity, with an average RMSF of 1.6 Å in MtCM^V^ compared to 2.5 Å (MtCM) for this region, the C-termini showed similarly high RMSF values in the two models (> 3 Å). Although Pro87 induced a kink at the C-terminus, this did not appear to affect the flexibility of the three terminal residues Asp88-Ala89-Met90.

### Kinetic Analysis to Probe Predicted Key Interactions of Engineered MtCM Variants

In the course of the directed evolution of MtCM^V^, the mutation L88D was only acquired after the H1-H2 loop-stabilizing substitutions T52P and V55D were already introduced. Guided by the outcome of the MD simulations, we therefore probed the kinetic impact of the innocuous single L88D exchange in the context of three different sets of MtCM variants to experimentally assess the benefit of the introduced negative charge for fine-tuning and optimizing catalytic efficiency. We looked at (*i*) mutating Asp88 in the MtCM^V^ sequence ^87^PDAM^90^ into Asn88 or Leu88, at (*ii*) directly introducing Asp88 into the MtCM wild-type sequence, as well as at (*iii*) the triple variant T52P V55D L88D. All variants were obtained in their native format, *i.e.*, with their native N terminus and without a His-tag, to allow for optimal comparison with the structural and computational results. The variants were purified by ion-exchange and size-exclusion chromatography from the *E. coli* host strain KA13, which is devoid of CM genes to rule out contamination by endogenous CMs (18,30). Subsequently, the enzymes’ kinetic parameters were characterized by a spectrophotometric chorismate depletion assay.

As shown in Table 2, removing the negative charge at residue 88 by replacing Asp with Asn in the top-evolved variant MtCM^V^ leads to a 2.5-fold drop in the catalytic efficiency *k*_cat_/*K*_m_ to 1.7 × 10^5^ M^−1^ s^−1^. This decrease is due both to a slightly lower catalytic rate constant (*k*_cat_) as well as a reduced substrate affinity (doubled *K*_m_). When residue 88 is further changed to the similarly sized but non-polar wild-type residue Leu88 in variant MtCM PLAM, the catalytic parameters essentially remain the same as for the Asn88 variant (Table 2), independently confirming the catalytic advantage of the negative charge introduced through Asp88.

**Table 2:** Catalytic CM activities of purified MtCM variants to experimentally address the importance of the Leu88 to Asp88 exchange that emerged the during directed evolution of MtCM^V^

| Variant | Mutations | $k_{\text{cat}}$ (s<sup>-1</sup>)<sup>a</sup> | $K_{\text{m}}$ (μM<sup>-1</sup>)<sup>a</sup> | $k_{\text{cat}}/K_{\text{m}}$ (M<sup>-1</sup> s<sup>-1</sup>)<sup>a</sup> |
| --- | --- | --- | --- | --- |
| MtCM<sup>V</sup> (PDAM) | PD/PDAM<sup>b</sup>, V62I, D72V, V11L, D15V, K40Q | 9.4 ± 1.3 | 22 ± 2 | 430,00 ± 30,000 |
| MtCM PNAM | MtCM<sup>V</sup>, D88N | 7.6 ± 0.2 | 45 ± 7 | 170,000 ± 20,000 |
| MtCM PLAM | MtCM<sup>V</sup>, D88L | 6.0 ± 0.4 | 38 ± 1 | 160,000 ± 10,000 |
| MtCM wt<sup>c</sup> | – | 1.7 ± 0.2 | 980 ± 80 | 1,700 ± 300 |
| MtCM L88D<sup>c</sup> | L88D | 3.0 ± 0.1 | 1110 ± 70 | 2,700 ± 300 |
| MtCM 3p3 | T52P V55D | 8.5 ± 0.0 | 450 ± 70 | 19,000 ± 3,000 |
| MtCM Triple | T52P V55D L88D | 11 ± 2 | 510 ± 130 | 22,000 ± 1,000 |

<sup>a</sup> All values are means from assays performed with at least two independently produced and purified protein batches with their standard deviations ( $\sigma_{n-1}$ ). The $k_{\text{cat}}/K_{\text{m}}$ parameters were obtained as the mean from averaging $k_{\text{cat}}/K_{\text{m}}$ values derived directly from individually fitted independent Michaelis-Menten plots with the error of the corresponding average.
<sup>b</sup> PD/PDAM indicates mutations T52P, V55D, R87P, L88D, G89A, and H90M.
<sup>c</sup> Whereas the default for measuring kinetics involved assays performed in 50 mM K-phosphate, pH 7.5, at 274 nm, the kinetic parameters of these low-performing variants were determined in 50 mM BTP, pH 7.5, at 310 nm. Measuring these variants in 50 mM K-phosphate, pH 7.5, resulted in ~40% reduction in $k_{\text{cat}}$ , as was already observed previously (21).

For the second set of variants that directly started out from the sluggish MtCM wild-type enzyme, a trend for an increase in catalytic activity upon the Leu88 to Asp88 replacement was observed (1.6-fold higher *k*_cat_/*K*_m_, reaching 2.7 × 10^3^ M^−1^ s^−1^; Table 2). This is mainly caused by an increase in *k*_cat_ rather than an altered substrate affinity. Interestingly, the L88D exchange together with T52P and V55D in the MtCM Triple variant does not lead to a significant increase in *k*_cat_/*K*_m_ compared to MtCM 3p3 (12), which just carries the two loop mutations T52P and V55D.

Thus, the substitution of Leu88 with Asp88 indeed results in a beneficial effect on the performance of MtCM. However, this effect is only prominent in combination with other selected mutations, such as those present in MtCM^V^. As a single mutation in the wild-type enzyme or on top of the H1-H2 loop mutations, the effect of L88D is less noticeable, if present at all.

In summary, a comparison of the dynamic behavior of wild-type MtCM in its apo and ligand-bound states with MtCM^V^ and MtCM^V55D^ revealed that the catalytically favorable conformation of the active site is achieved by the interplay of several interactions, balancing charges and entropic disorder of the H1-H2 loop. Structuring is promoted, in particular, by increasing the number of the negatively charged carboxylate groups that can both shield the electrostatic charge of the various arginine side chains within or next to the active site and orient catalytically important residues by hydrogen bonding and salt bridge formation. Simulations of MtCM^V^ revealed the special importance of Asp55 in the V55D variant for coordinating Arg18’ and Arg46, thus promoting the preorganization of the active site region. These results echo the conclusions from directed evolution, which also identified the V55D substitution as the most important contributor for catalytic enhancement, causing a 12-fold increase in *k*_cat_/*K*_m_ (28). At the same time, we determined and rationalized the more subtle and context-dependent effect of the L88D exchange that introduced an additional negative charge for electrostatic preorganization of the active site. Overall, the high catalytic activity of MtCM^V^ clearly results from many individual larger and smaller contributions mediated by substitutions at diverse locations within the enzyme structure.

## DISCUSSION

### Important Activating Factors in MtCM^DS^ and MtCM^V^

MtCM has intrinsically low activity but can be activated through formation of a heterooctameric complex with MtDS, which aligns crucial active site residues to catalytically competent conformations. Most importantly, binding to MtDS induces pre-organization of Arg46 into a catalytically favorable conformation (Fig. 2B), *via* H-bonding to the carbonyl oxygens of Thr52 and Arg53 (21). Arg46 is the crucial catalytic residue interacting with the ether oxygen of Bartlett’s transition state analog (TSA) (51) in the complex with MtDS (PDB ID: 2W1A (21)) (Figs. 1B/F and 2B); upon mutation to Lys, the enzyme’s efficiency drops 50-fold (21).

Both MtCM^DS^ and MtCM^V^ exhibit a kinked H1-H2 loop conformation (Figs. 1C/D and 2A), which was hypothesized to be important for increased catalytic efficiency (12). However, in MtCM^V^ and MtCM^V55D^, the kink is exacerbated by crystal contacts, which are different in the two crystal forms (Fig. S2). This kink is much less prominent in wild-type MtCM, or even MtCM^T52P^ (Figs. 2A and S1B), and completely lost during the simulations of MtCM^WT^ (we did not carry out simulations on the single variant MtCM^T52P^). Thus, this conformation may well be a crystallization artifact rather than a prerequisite for an active MtCM.

Nevertheless, pre-organization and pre-stabilization appear to be of crucial importance for the catalytic prowess of MtCM. V55D is the single substitution among the MtCM^V^ mutations that resulted in the largest boost in catalytic activity (12-fold enhancement) (12). This residue is located on the C-terminal side of the H1-H2 loop (Fig. 1D/E) and forms a salt bridge to the catalytically important Arg46 at the top of helix H1 (Fig. 5A), an interaction that is also observed in the crystal structure of MtCM^V55D^ (Fig. S1G/H). During the MD simulations of MtCM^V^ and the single variant MtCM^V55D^, the presence of Asp55 reduced the mobility of active site residues. By interacting with Arg18’ and Arg46, this residue helps to pre-organize the active site for catalysis and reduce unfavorable conformational fluctuations caused by electrostatic repulsion in the absence of a substrate. This is supported by the lower RMSF values of MtCM^V^ compared to uncomplexed wild-type MtCM (Table 1) and by a slightly higher melting temperature of MtCM^V55D^ (ΔT = 3 °C from preliminary differential scanning fluorimetry (DSF) measurements; preliminary data). By decreasing thermal fluctuations in the active site, Asp55 likely also reduces the entropic penalty associated with substrate binding. Pro52 appears to exert a similar stabilizing effect on the protein, despite the rather small structural changes, as suggested by a 2 °C increase in melting temperature of MtCM^T52P^ in DSF experiments compared to MtCM (preliminary data). This single substitution raises the *k*_cat_/*K*_m_ of the enzyme by a factor of six (12). It is worth noting that the simultaneous substitution of T52P and V55D increased the melting temperature by 6 °C (monitored by circular dichroism spectroscopy) and boosted *k*_cat_/*K*_m_ by 22-fold (12).

### Importance of the C-Terminus

MtCM activation by MtDS involves a change in conformation of the C-terminus of MtCM and its active site H1-H2 loop (21). Specifically, a salt bridge is formed between the C-terminal carboxylate of MtCM (which is repositioned upon MtDS binding) and loop residue Arg53, possibly bolstered by a newly formed hydrophobic interaction between Leu88 and Leu54 (Fig. S3A). The 1-µs simulations suggest that salt-bridge formation with Arg53 occurs in solution in all tested cases, whereas the hydrophobic contact is less important.

Directed evolution experiments carried out by randomizing the final four C-terminal positions 87-90 of MtCM had previously revealed that a great variety of residues with quite distinct physical-chemical properties are compatible with a functional catalytic machinery (26). Conserved positions emerged only when probing for an intact activation mechanism by MtDS (26). Still, when residues 87-90 of MtCM^V^ were evolved from Arg-Leu-Gly-His to Pro-Asp-Ala-Met (Fig. 1E), an increase in *k*_cat_/*K*_m_ by roughly a factor of four was achieved (12). Here, we resolved this apparent paradox by investigating C-terminal factors important for the fine-tuned optimization of CM function. Even though the replacement R87P induced a kink in the structure, the presence of the proline did not appear to have a major influence in the simulations. Notably, the C-terminal substitutions together result in a change in net charge from +1 to −2, including the terminal carboxylate, providing the basis for more extensive electrostatic interactions with the positively charged Arg53 than is possible for wild-type MtCM. Indeed, our kinetic analysis of Asp88-containing MtCM variants demonstrates that this residue increases CM’s catalytic efficiency (Table 2). The fact that Asp88 did not significantly augment *k*_cat_/*K*_m_ in the context of the MtCM double variant T52P V55D (*i.e.*, MtCM Triple; Table 2) suggests that the extent of catalytic improvement by L88D depends on the particular structural context.

Our simulations indicate that in free wild-type MtCM, an interaction of the C-terminal carboxylate with the key active site residue Arg46 is possible, but infrequent due to fluctuations (Fig. 6A; Table 1). In contrast, in MtCM^V^ and MtCM^DS^ the side chain of Arg46 points towards the catalytic pocket (Figs. 5 and 2B), and any unproductive reorientation of Arg46 towards the C-terminus would easily result in a clash with the H1-H2 loop. Thus, an additional feature of this loop may be to act as a conditional shield (illustrated for MtCM^V^ in Fig. 7). In the conformation assumed in MtCM^V^ and MtCM^DS^, this loop blocks reorientation of Arg46 towards the C-terminus and hence prevents an unproductive conformation accessible for free wild-type MtCM. MtCM^DS^ and MtCM^V^ use different means to correctly position active site residues, which correlates with a bent H1-H2 loop in both cases. This is either achieved through conformational changes imposed upon MtCM^DS^ by MtDS binding, or by establishing a salt bridge across the active site, between Arg46 and Asp55, as seen for MtCM^V^ and also for the single variant MtCM^V55D^ (Fig. 5 and S1E/G/H).

**Figure 7.**
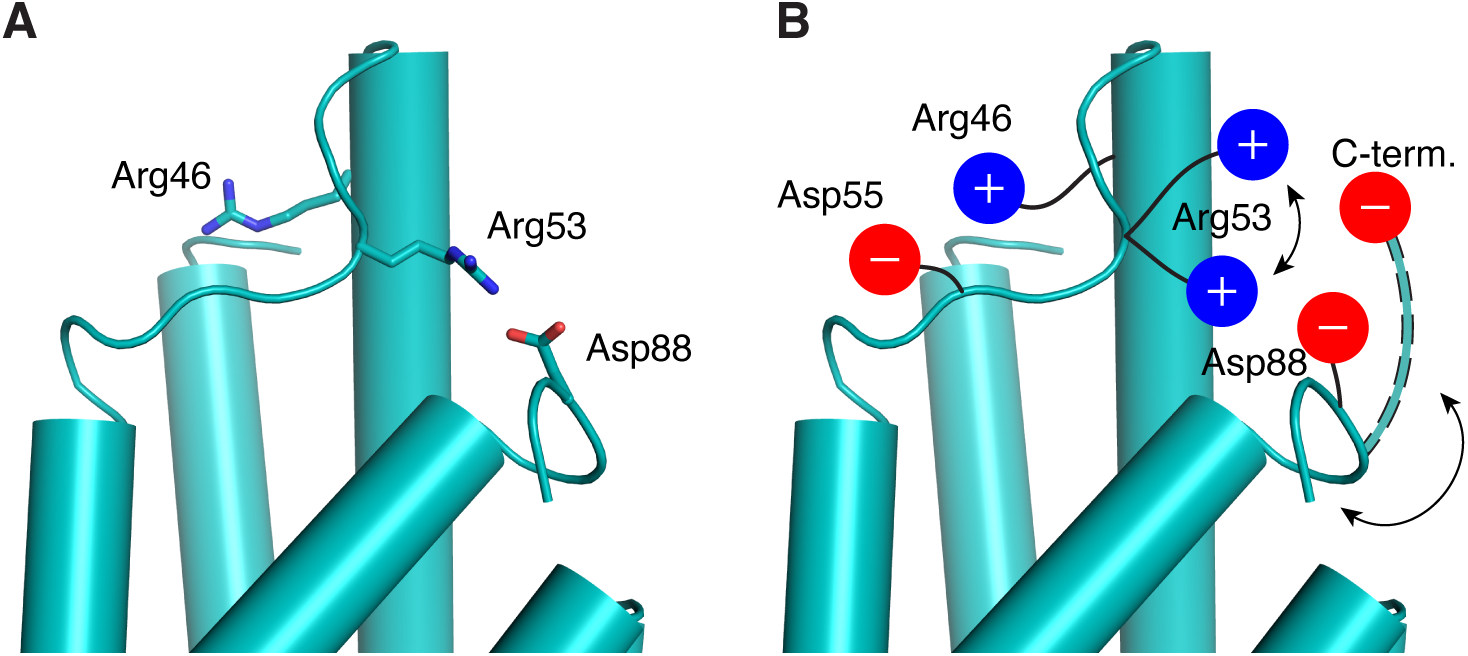
Shielding interaction mediated by the H1-H2 loop. (**A**) Conformation of important Arg residues in chain A of MtCM^V^ (cyan) after 31.7 ns of MD simulations. The key active site residue Arg46 is positioned on the opposite side of the H1-H2 loop, which in turn is bolted to the C-terminus by a salt bridge between Arg53 and Asp88 (cartoon representation, with side chains shown as sticks). (**B**) Cartoon summarizing the important stabilizing interactions in the top-evolved variant MtCM^V^ depicted in (A) that properly position Arg46 for catalysis. Asp55 stabilizes the stretched-out conformation of Arg46, whereas alternating salt bridges accessible for Arg53 with the negatively charged groups present in the C-terminal region hinder Arg46 from adopting an unfavorable interaction with the C-terminal carboxylate. One example of an alternative backbone conformation that allows for interactions between the C-terminal carboxylate and Arg53 is depicted with dashed outlining.

### General Implications for CM Catalysis

It is obviously impossible to directly transfer our findings of critical detailed molecular contacts from the AroQ_δ_ subclass CM of *M. tuberculosis* to the evolutionary distinct AroH class CMs, or even to the structurally and functionally divergent AroQ_α_, AroQ_β_, and AroQ_γ_ subclasses (53). Neither of those groups of CMs have evolved to be deliberately poor catalysts that become proficient upon regulatory interaction with a partner protein such as MtDS (21). To be amenable to ‘inter-enzyme allosteric’ regulation (28), the H1-H2 loop in MtCM must be malleable and allow for conformational switching between a poorly and a highly active form. In contrast, this region is rigidified in a catalytically competent conformation in the overwhelming majority of CMs from other subclasses. This is exemplified by the prototypic EcCM (AroQ_α_ subclass) and the secreted *MtCM (AroQ_γ_), which possess the sequence ^45^**P**VR**D**^48^ and ^66^**P**IE**D**^69^, respectively, at the position corresponding to the malleable H1-H2 loop sequence ^52^**T**RL**V**^55^ of wild-type MtCM (12). Remarkably, the two most impactful substitutions T52P and V55D occurring during the evolution of MtCM^V^ have led to the tetrapeptide sequence ^52^**P**RL**D**^55^, with both Pro and Asp being conserved in naturally highly active CMs (12).

The AroQ_δ_ subclass CM from *Corynebacterium glutamicum* is another structurally well characterized poorly active CM (*k*_cat_/*K*_m_ = 110 M^−1^ s^−1^) that requires complex formation with its cognate DAHP synthase for an impressive 180-fold boost in catalytic efficiency (54). In that case, inter-enzyme allosteric regulation involves a conformational change of a different malleable segment between helices H1 and H2. Thus, while the molecular details important for the activation of a particular AroQ_δ_ CM cannot be transferred directly from one system to another, our findings suggest as a general regulatory principle the deliberate and reversible destabilization of a catalytically critical loop conformation.

In both the *M. tuberculosis* (12) and the *C. glutamicum* systems (54) crystal contacts in the H1-H2 loop region impede the structural interpretation of the activity switching. The MD simulations shown here represent an interesting alternative approach to dynamic high-resolution structure determination methods for sampling the conformational space adopted by malleable peptide segments with and without ligands.

## CONCLUSIONS

MD greatly aided the analysis of crystal structures that were compromised or biased by extensive crystal contacts at the most interesting structural sites. Our aim was to obtain insight into the crucial factors underlying CM activity by comparing the structure and dynamics of the poorly active wild-type MtCM (*k*_cat_/*K*_m_ = 1.7 × 10^3^ M^−1^ s^−1^) with a highly efficient MtCM variant (MtCM^V^; *k*_cat_/*K*_m_ = 4.3 × 10^5^ M^−1^ s^−1^), which emerged from directed evolution experiments. Both in MtDS-activated wild-type MtCM and in MtCM^V^, high activity correlated with a kinked H1-H2 loop conformation and an interaction of this region with the C-terminus of MtCM. The autonomously fully active variant MtCM^V^ had mutations in both of these regions that augment these structural features. In this report we focussed on substitutions T52P, V55D, and L88D.

The active site of all natural CMs contains a high density of positive charges. In MtCM, four arginine residues (Arg18’, Arg35, Arg46, and Arg58, of which Arg18’ is contributed by a different MtCM protomer) are responsible for binding and rearranging the doubly negatively charged substrate chorismate. Only one of these residues (Arg35) is firmly in position before the substrate enters the active site. Of critical importance for catalysis is Arg46. During the MD simulations, Arg46 competes with another arginine residue (Arg53) for binding to the C-terminal carboxylate (Fig. 6A) and adopts a catalytically unproductive conformation unless an aspartate residue (Asp55 or Asp88) comes to its rescue. As shown here, Asp55 not only properly orients Arg46 for catalysis, but additionally stabilizes the active site. Together with T52P, which pre-orders the H1-H2 loop, the V55D exchange results in reduced mobility of residues in the active site through stabilizing interactions, thereby pre-organizing it for efficient catalysis and lowering the entropic cost of substrate binding. Another aspartate residue (Asp88), also acquired in the top-evolved MtCM^V^ (12), helps to balance charges, and—by interacting with Arg53—imposes a steric block that prevents non-optimal positioning of Arg46 (Fig. 7), explaining why the L88D exchange can increase *k*_cat_/*K*_m_ by about 2-3 fold.

In summary, we tested our hypotheses on the specific importance of critical substitutions acquired during the directed evolution of MtCM^V^, namely T52P, V55D, and L88D by investigating single variants as well as combinations with other residue replacements that were found to augment catalysis. The variants were characterized by crystallography, MD simulations, and enzyme kinetics. The two residues Pro52 and Asp55 exert a major impact by pre-stabilization and pre-organization of catalytically competent conformations of active site residues, while Asp88 fine-tunes and optimizes the catalytic process. By expanding on the previous directed evolution studies, we have shown here how the accumulated set of amino acid substitutions found in MtCM^V^ has resulted in an activity level matching that of the most active CMs known to date (12).

## SUPPORTING INFORMATION

This article contains supporting information (Supporting Figures S1-S4 and Supporting Table S1).

## ACKNOWLEDGEMENTS

We would like to thank Regula Grüninger-Stössel for help with the construction of the T52P and V55D variants of MtCM, and Joel B. Heim for collecting X-ray data for MtCM^T52P^. We would further like to acknowledge the European Synchrotron Radiation Facility for provision of synchrotron radiation facilities and thank Montserrat Soler Lopez and Daniele De Sanctis for assistance in using beamlines ID30A-3/MASSIF-3 and ID29, respectively. Finally, we acknowledge services provided by the MoBiAS facility, Laboratory of Organic Chemistry, ETH Zurich, and by the UiO Structural Biology Core Facilities.

## AUTHOR CONTRIBUTIONS

U.K. conceived the study. H.V.T., P.K. and Mi.C. were additionally involved in the planning of the experiments. H.V.T. performed most of the calculations, transformed, produced, purified and crystallized the two single MtCM variants, and solved the crystal structure of MtCM^V55D^, supervised by Mi.C. and U.K., respectively. Ma.C. contributed with additional simulations, supervised by Mi.C. T.K. solved the crystal structure of MtCM^T52P^ and refined the crystal structures of both MtCM variants, supervised by G.C. and U.K., who also validated the structures. L.B. constructed, produced and purified additional sets of MtCM variants and characterized their kinetic parameters to validate computational results, and K.W.-R. designed and constructed the MtCM variants T52P and V55D and prepared the final figures; both were supervised by P.K. The initial version of the manuscript was written by H.V.T. and U.K., which was complemented with contributions from all authors and revised by P.K., Mi.C. and U.K.

