## Supplementary Information for "What drives chorismate mutase to top performance? Insights from a combined *in silico* and *in vitro* study"

### LIST OF MATERIAL INCLUDED:

**Figures S1-S4** (S1, Crystal structures and electron density; S2, Crystal contacts; S3-4, MD snapshots)

**Table S1** (Data collection and refinement statistics)

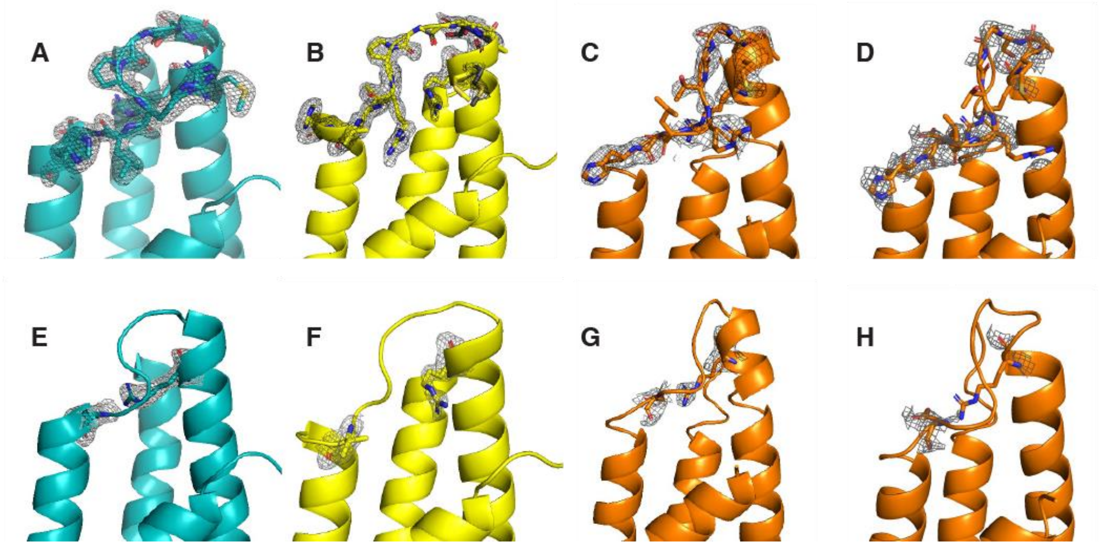


**Figure S1. Crystal structures and electron density maps shown for catalytically important regions.** Top row, active site H1-H2 loop residues ($^{47}$MASGGPRLVHS$^{57}$). Bottom row, residue 55 and the catalytically crucial Arg46. **(A)** and **(E)**, top-evolved MtCM$^{V}$ (PDB ID: 5MPV (12); cyan); **(B)** and **(F)**, MtCM$^{T52P}$ (PDB ID: 6YGT, this work, with H1-H2 loop modeled in two alternative conformations; yellow); **(C)** and **(G)**, MtCM$^{V55D}$, protomer A (this work; orange); **(D)** and **(H)**, MtCM$^{V55D}$, protomer B (this work; orange). The MtCM$^{V55D}$ structure is of low quality, precluding final refinement; therefore, the coordinates were not deposited in the PDB. All electron density maps are $\sigma_A$-weighted 2m*F*o-D*F*c maps, depicted at σ =1.0.

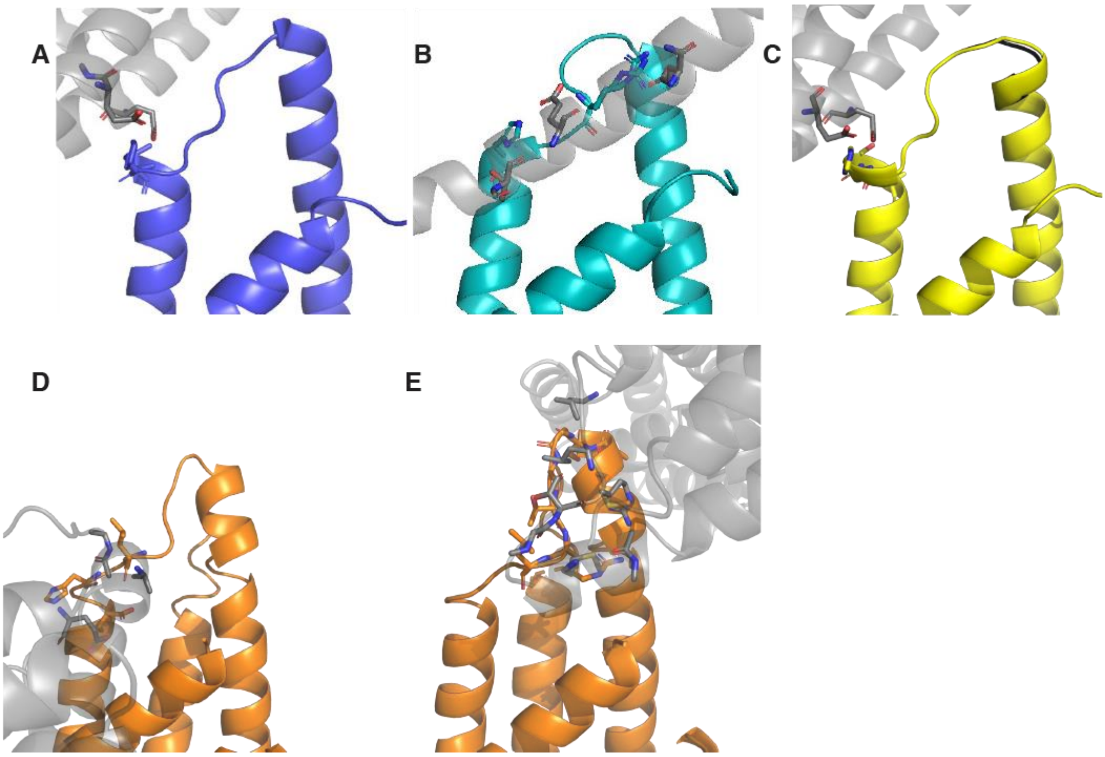


**Figure S2. Crystal contacts of the H1-H2 loop with the closest neighboring molecule (shown in grey). (A)** Wild-type MtCM (PDB ID: 2VKL (21); purple), **(B)** Top-evolved MtCM$^{V}$ (PDB ID: 5MPV (12); cyan), **(C)** MtCM$^{T52P}$ (PDB ID: 6YGT, this work; yellow), **(D)** MtCM$^{V55D}$, protomer A (this work; orange), **(E)** MtCM$^{V55D}$, protomer B (this work; orange). The structures in panels A and C have the same crystal form.

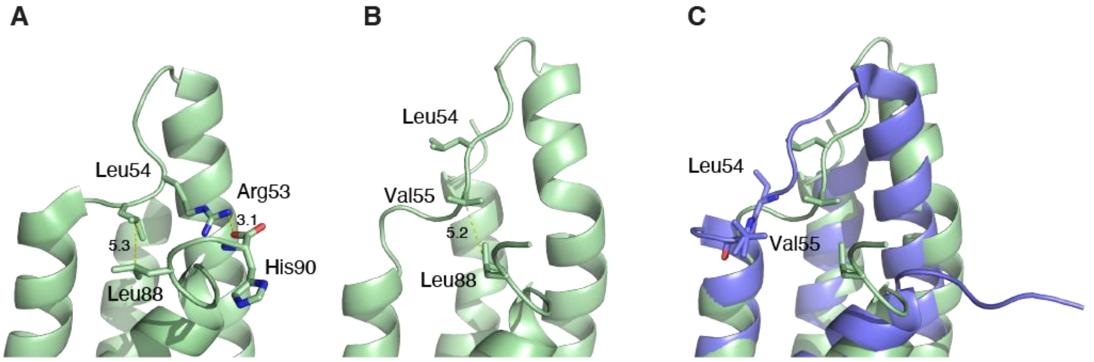


**Figure S3. Interactions in MtCM between its C-terminus and H1-H2 loop.** (**A**) Cartoon illustration of MtCM$^{DS}$ (MtCM from the MtCM-MtDS crystal structure; PDB ID: 2W19 (20)). MtCM$^{DS}$ is colored green and residues involved in prominent interactions are shown as sticks. Two interactions are highlighted: hydrophobic contacts between Leu54 of the H1-H2 loop and Leu88 close to the C-terminus of MtCM (distance measured between $C_\gamma$ of Leu side chains), and a salt bridge between Arg53 of the H1-H2 loop and the C-terminal carboxylate (His90). (**B**) Cartoon illustration of van der Waals interactions observed for chain A of MtCM$^{DS}$ at time step 250 (25 ns simulation). Note the distinct conformation of the H1-H2 loop, with Val55 temporarily taking over the role of Leu54. However, this conformational change is catalytically unfavorable, as it interferes with substrate binding to the Val55 main chain amide group, as shown in Fig. 1F. (**C**) Superimposition of MtCM$^{DS}$ (simulated structure from **B**, green) with MtCM crystal structure (purple, PDB ID: 2VKL (20)). The corresponding residues Val55 and Leu54 in the two structures, shown as sticks, occupy shifted positions, but in similar orientations in the respective H1-H2 loops.

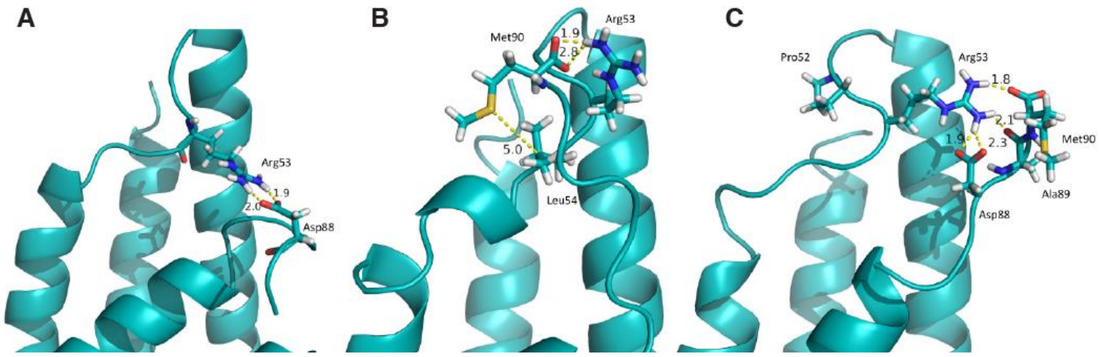


**Figure S4. MD snapshots of interactions between the C-terminus of MtCM$^{V}$ and its H1-H2 loop.** The C-terminus is very flexible, but recurring interactions are made by carboxylic acid groups from the C-terminal sequence (C-terminal carboxylate or Asp88 side chain) to Arg53 of the H1-H2 loop. Representative frames are shown in cartoon representation, with relevant residues as sticks. (**A**) Chain A at 15.1 ns, (**B**) Chain B at 17.8 ns, (**C**) Chain A at 4.1 ns (intermediate conformation).

### Table S1 – Data collection and refinement statistics

| | MtCM$^{T52P}$ | MtCM$^{V55D}$ |
|---|---|---|
| ***Data collection*** | | |
| Beamline | ESRF ID3-A30/MASSIF-3 | ESRF ID29 |
| Wavelength (Å) | 0.9677 | 0.9753 |
| Space group | $P4_32_12$ | $P2\ 2_1\ 2_1$ |
| Cell parameters - *a*, *b*, *c* (Å) | 59.6, 59.6, 46.6 | 32.2, 59.7, 72.1 |
| Protein chains in a.s.u. | 1 | 2 |
| Matthew's coefficient (Å$^3$/Da) | 2.0 | 1.7 |
| Resolution (Å)[a] | 36.7-1.64 (1.72-1.64) | 45.94-2.06 (2.25-2.06) |
| $CC_{1/2}$ (%)[a, b] | 99.5 (45.1) | 99.9 (47.6) |
| Mean I/σ(I)[a] | 15.7 (1.1) | 14.6 (1.1) |
| Completeness (%)[a] | 88.1 (33.4) | 82.4 (22.3) |
| Number of unique reflections [a] | 9570 (479) | 7445 (438) |
| Multiplicity[a] | 8.7 (9.8) | 6.0 (6.1) |
| Wilson B-factor (Å$^2$) | 22.2 | 57.8 |
| ***Refinement***[d] | | |
| Resolution range (Å) | 29.8-1.64 | |
| $R_{work}/R_{free}$ (%)[c] | 24.0/26.5 | |
| Average *B*-factor (Å$^2$) | 32.6 | |
| Number of atoms | | |
| Protein | 674 | |
| Water | 18 | |
| r.m.s.d. from ideal geometry | | |
| Bond lengths (Å) | 0.01 | |
| Bond angles (deg.) | 1.2 | |
| Ramachandran plot[e] | | |
| Favored (%) | 93.1 | |
| Allowed (%) | 5.2 | |
| Outliers (%) | 0.0 | |
| PDB code | 6YGT | |

[a] Values in parentheses refer to highest resolution shell. The data completeness falls below the 95% threshold beyond ~1.85 Å and ~2.40 Å resolution for MtCM$^{T52P}$ and MtCM$^{V55D}$, respectively, which can be considered the effective resolution of the data sets.

[b] Reflections up to the highest resolution limit were included assessing the data using the $CC_{1/2}$ parameter, as suggested by Diederichs and Karplus (55,56).

[c] $R_{free}$ was calculated from 5% of randomly selected reflections for each data set.

[d] Refinement of MtCM$^{V55D}$ stalled at $R_{work}/R_{free}$ values of 27.6/34.9%. The structure was therefore not included in this Table or submitted to the PDB.

[e] Calculated with *SFCHECK* (55). Ser49 (and Gly51) have torsion angles bordering to outlier regions in the Ramachandran plot, and were identified as outliers by the PDB.
