## Supplementary figures and images for "What drives chorismate mutase to top performance? Insights from a combined *in silico* and *in vitro* study"

### Supplementary Figure 1

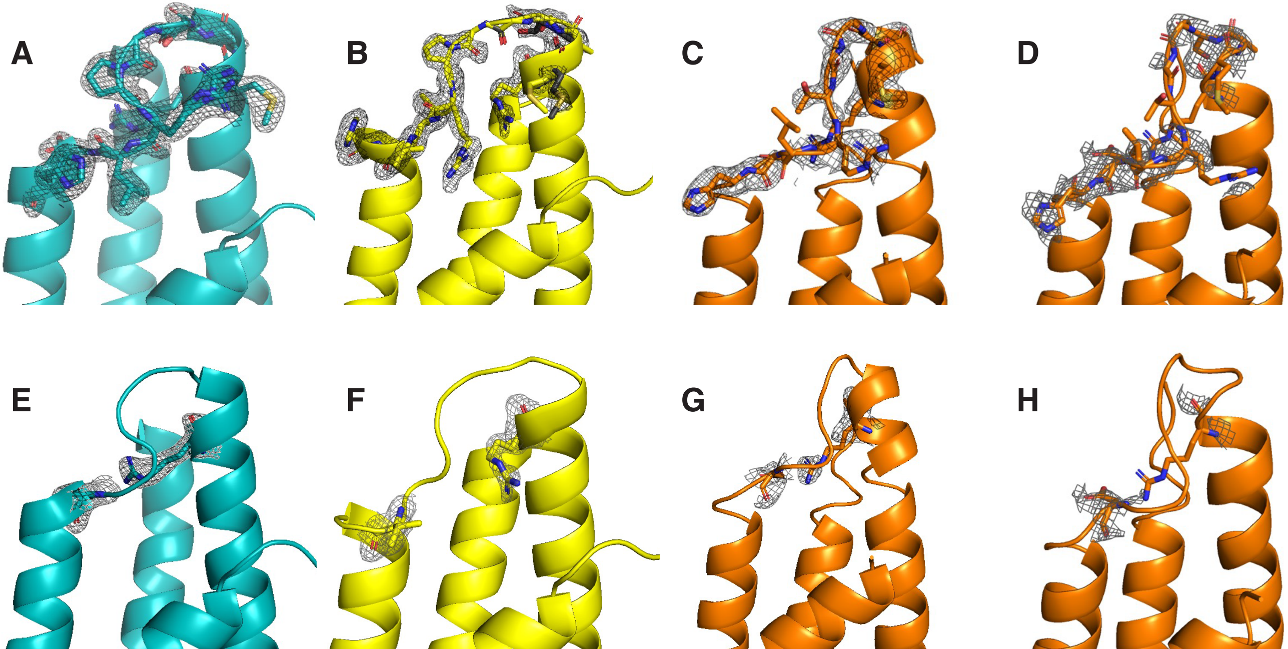

### Supplementary Figure 2

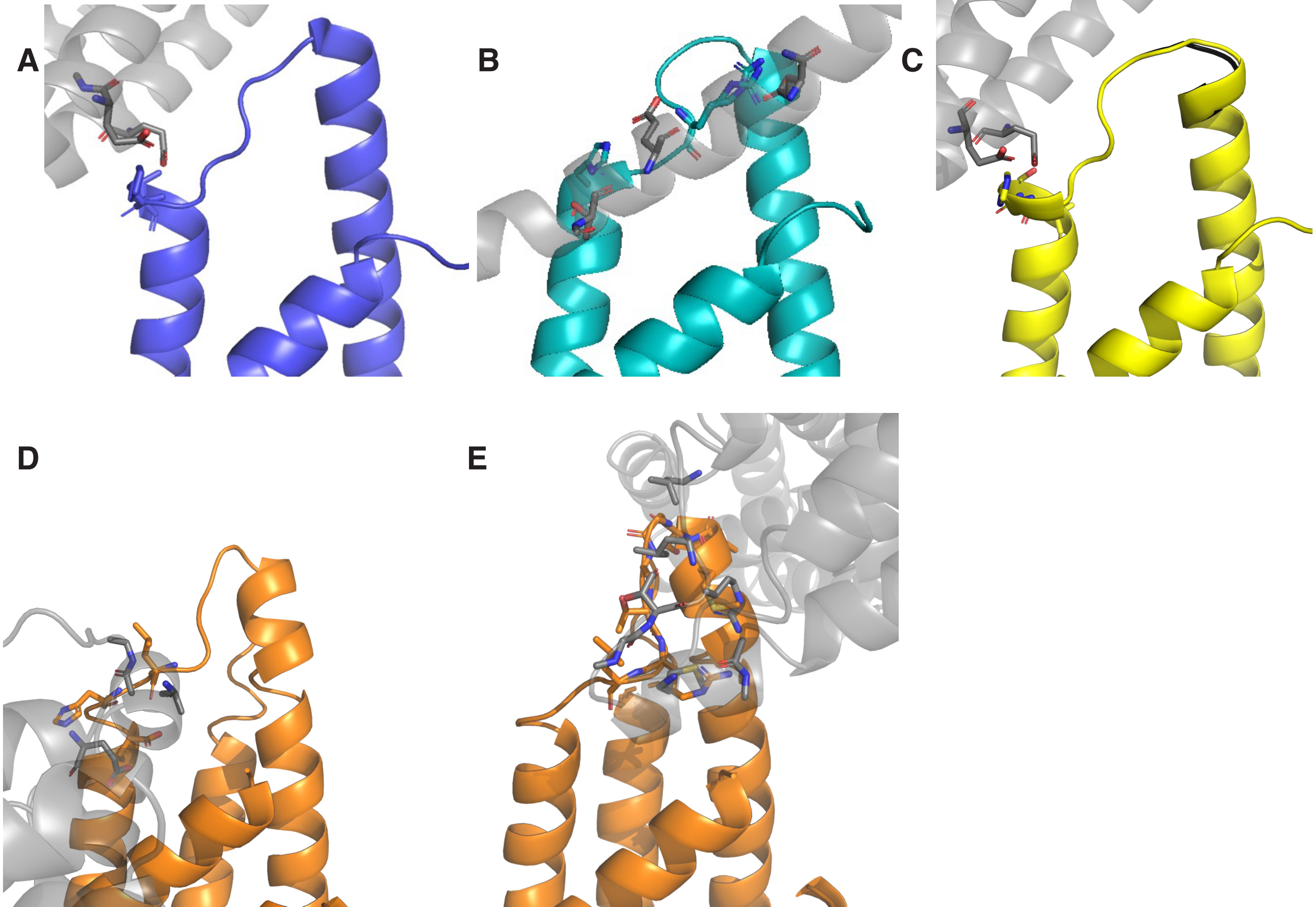

### Supplementary Figure 3

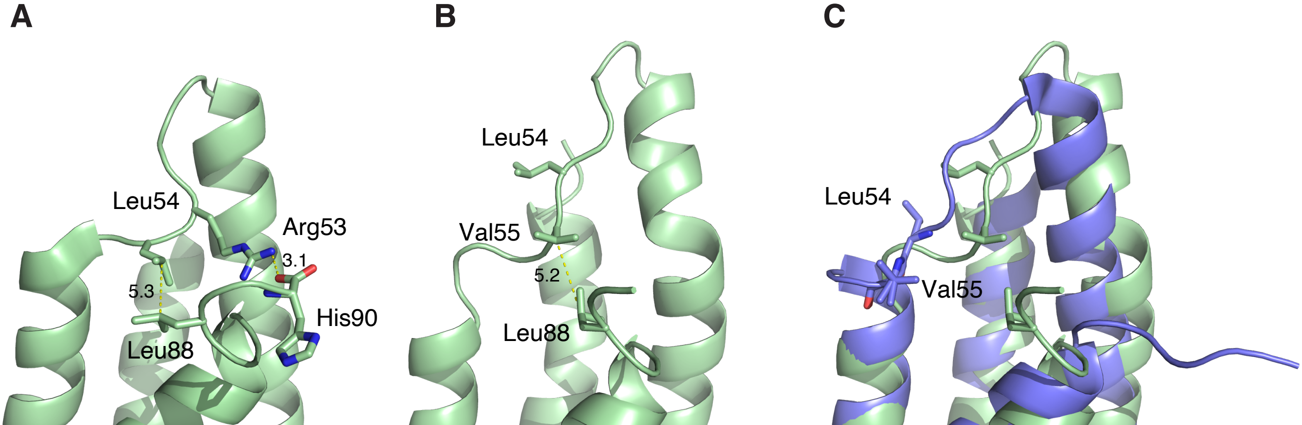

### Supplementary Figure 4

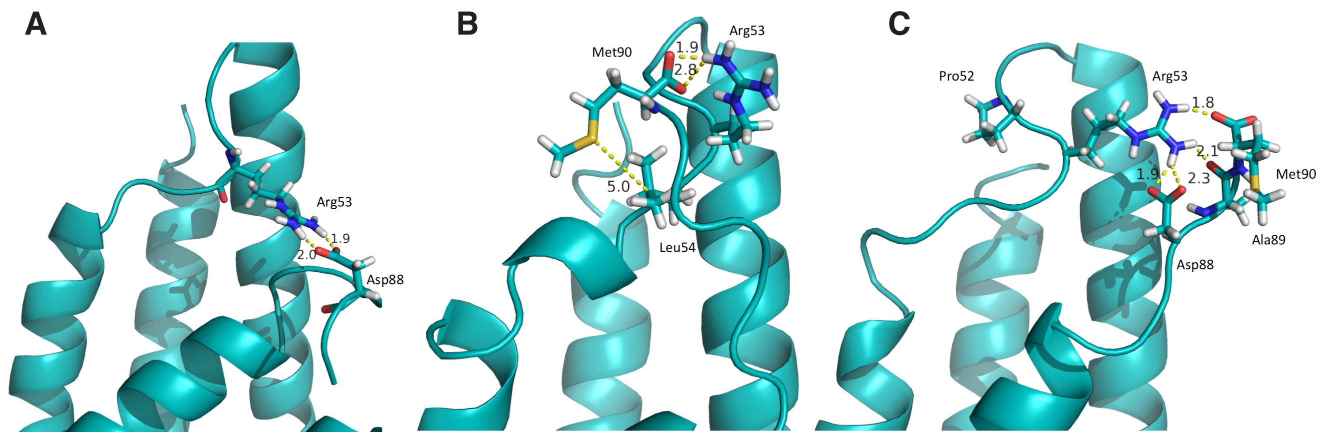
